# Mechano-redox control of Mac-1 de-adhesion from ICAM-1 by protein disulfide isomerase promotes directional movement of neutrophils under flow

**DOI:** 10.1101/2022.03.29.486223

**Authors:** Alexander Dupuy, Camilo Aponte Santamaría, Adva Yeheskel, Frauke Gräter, Philip J. Hogg, Freda H. Passam, Joyce Chiu

## Abstract

Macrophage-1 antigen or Mac-1 (CD11b/CD18, αMβ2) is a leukocyte integrin essential for firm adhesion of neutrophils, lymphocytes and monocytes against flow when recruited to the endothelium. To migrate to the site of inflammation, leukocytes require coordinated adhesion and de-adhesion for directional movement. The vascular thiol isomerase, protein disulfide isomerase (PDI), was found by fluorescence microscopy to colocalize with high affinity Mac-1 at the trailing edge of stimulated neutrophils when adhered to ICAM-1 under fluid shear. From differential cysteine alkylation and mass spectrometry studies, PDI cleaves two allosteric disulfide bonds, C169-C176 and C224-C264, in the βI domain of the β2 subunit, and in mutagenesis and cell transfection studies, cleavage of the C224-C264 disulfide bond was shown to selectively control Mac-1 dis-engagement from ICAM-1 under fluid shear. Molecular dynamics simulations and binding of conformation-specific antibodies reveal that cleavage of the C224-C264 bond induces conformational change and mechanical stress in the βI domain that allosterically alters exposure of an αI domain epitope and shifts Mac-1 to a lower affinity state. From studies of neutrophil adherence to ICAM-1 under fluid shear, these molecular events promote neutrophil motility in the direction of flow at high shear stress. In summary, shear-dependent PDI cleavage of neutrophil Mac-1 C224-C264 disulfide bond triggers Mac-1 de-adherence from ICAM-1 at the trailing edge of the cell and enables directional movement of neutrophils during inflammation.

## Introduction

The integrin macrophage-1 antigen or Mac-1 (CD11b/CD18, αMβ2) is essential for the recruitment of leukocytes to sites of infection or injury. It binds to a variety of ligands including intercellular adhesion molecule 1 (ICAM-1), fibrinogen, complement fragment iC3b and CD40 ligand (CD40L) to elicit an inflammatory response. To migrate to the site of infection and injury, circulating leukocytes tether and roll on vessel wall by interacting with selectins expressed on endothelial cells, reducing their velocity. Their initial contacts trigger G-protein coupled receptors for inside-out signaling leading to integrin activation and binding to endothelial ICAM-1 (Ley et al., 2007). Integrin-mediated adhesion leads to assembly of focal adhesions and cytoskeletal remodeling to enable cell spreading and firm anchorage of leukocytes against shear force (Lefort et al., 2009; Smith et al., 2005; Takami et al., 2001). As leukocytes become polarized for directional crawling, their trailing edge needs to detach to move forward. Various mechanisms have been reported to enable cell de-adhesion. For instance, shedding of integrin by metalloproteinases has been described to enable exit of macrophages from the site of inflammation (Gomez et al., 2012). Internalization of integrin by clathrin-mediated endocytosis is another important mechanism that allows disassembly of focal adhesions and detachment of cells from substrata (Bai et al., 2017; Ezratty et al., 2009). These mechanisms rely on removal of functional integrin from the cell. We have described a mechanism of integrin dis-engagement that involves allosteric changes in the integrin binding sites (Passam et al., 2018).

De-adhesion of platelet integrin αIIbβ3 (GPIIb/IIIa, CD61/CD41) from fibrinogen occurs via force-dependent cleavage of an allosteric disulfide bond in the integrin binding site (Passam et al., 2018). A member of the vascular thiol isomerase family, ERp5, cleaves a disulfide bond in the β3 subunit to release platelets from fibrinogen. The archetypal thiol isomerase, protein disulfide isomerase (PDI), has been demonstrated to be essential in Mac-1-dependent neutrophil migration during inflammation. Mac-1 becomes upregulated during inflammation to mediate neutrophil adhesion and crawling (Sumagin et al., 2010). Conditional knockout of PDI in murine neutrophils led to their impaired adhesion and crawling on inflamed endothelium (Hahm et al., 2013).

Here, we report a mechano-redox mechanism that mediates Mac-1 de-adhesion selectively from ICAM-1. PDI colocalizes with high affinity Mac-1 at the trailing edge of neutrophils, and cleaves a disulfide bond in the headpiece of the β2 subunit that changes Mac-1 conformation to a lower affinity state. PDI cleavage of the Mac-1 disulfide bond promotes neutrophil movement in the direction of fluid shear. This mechano-redox regulation by PDI provides a mechanism for neutrophils to de-adhere from ICAM-1 that is essential for directional movement and migration during inflammation.

## Results

### Surface PDI colocalizes with high affinity Mac-1 at the trailing edge of neutrophils adhering to ICAM-1

Hahm et al. demonstrated that PDI-null neutrophils exhibited defective migration on TNFα inflamed endothelium and that such migration could be restored by addition of recombinant PDI (Hahm et al., 2013). As neutrophil adhesion and crawling on endothelium is predominantly dependent on Mac-1 binding to endothelial ICAM-1, co-localization of surface PDI and Mac-1 was measured in fixed neutrophils adhered to immobilized ICAM-1. Using anti-PDI antibody DL-11, low level of PDI was detected on the surface of untreated neutrophils adhered to ICAM-1 (**Figure 1 – figure supplement 1**). Using anti-CD11b antibody, CBRM1/5, that recognizes an activation-specific epitope in the I domain of αM (αI domain) that is exposed only in Mac-1 at high affinity state (Oxvig et al., 1999), high affinity Mac-1 was hardly detected in resting neutrophils. Upon stimulation with fMLF, PDI and high affinity Mac-1 were readily detected on the neutrophil surface. Increased cell surface PDI was detected in fMLF-stimulated neutrophils in accordance with a previous report (Hahm et al., 2013), which was accompanied by Mac-1 upregulation (Kishimoto et al., 1989). PDI predominately colocalized with high affinity Mac-1 clusters associated with focal adhesion points for firm adhesion of neutrophils on ICAM-1.

To determine if surface PDI colocalizes with high affinity Mac-1 during neutrophil crawling, neutrophils were stained with anti-PDI antibody and CBRM1/5, stimulated by fMLF and perfused onto microfluidic chips coated with ICAM-1 and left to settle. Adhered neutrophils were then subjected to shear stress representing venous or arterial vessel at 0.7 or 5.6 dynes/cm^2^ (Sakariassen et al., 2015), respectively, and imaged in real-time by confocal microscopy.

Adhered neutrophils exhibited polarized morphology in presence of fluid shear with leading and trailing edges clearly defined on DIC images (Valignat et al., 2014). PDI was found to localize in the trailing edge of neutrophils while high affinity Mac-1 also predominately formed clusters in the trailing edge but was also detected in the middle of crawling neutrophils as previously reported (Hyun et al., 2019) (**Figure 1A**). The fractions of PDI and Mac-1 that overlapped in leading and trailing edges were measured and expressed as Manders’ colocalization coefficients (**Figure 1B** and **Supplementary File 1 Table S1**). Half of PDI fluorescence at the trailing edge was found to overlap with Mac-1 fluorescence at the trailing edge (Manders’ colocalization coefficient for PDI is 0.5512 ± 0.1957 and 0.5287 ± 0.2368 at 0.7 and 5.6 dynes/cm^2^, respectively) as compared to only 13-20% of PDI fluorescence in the leading edge overlapping with Mac-1 fluorescence in the leading edge (Manders’ colocalization coefficient for PDI: 0.1969 ± 0.1355 and 0.132 ± 0.1257 at 0.7 and 5.6 dynes/cm^2^, respectively). Similarly, half of Mac-1 fluorescence at trailing edge was found to overlap with PDI fluorescence at the trailing edge as compared to only 20-25% of Mac-1 fluorescence in the leading edge (**Supplementary File 1 Table S1**). This indicates that PDI and high affinity Mac-were more colocalized in the trailing edge than in the leading edge of crawling neutrophils.

**Figure 1.**
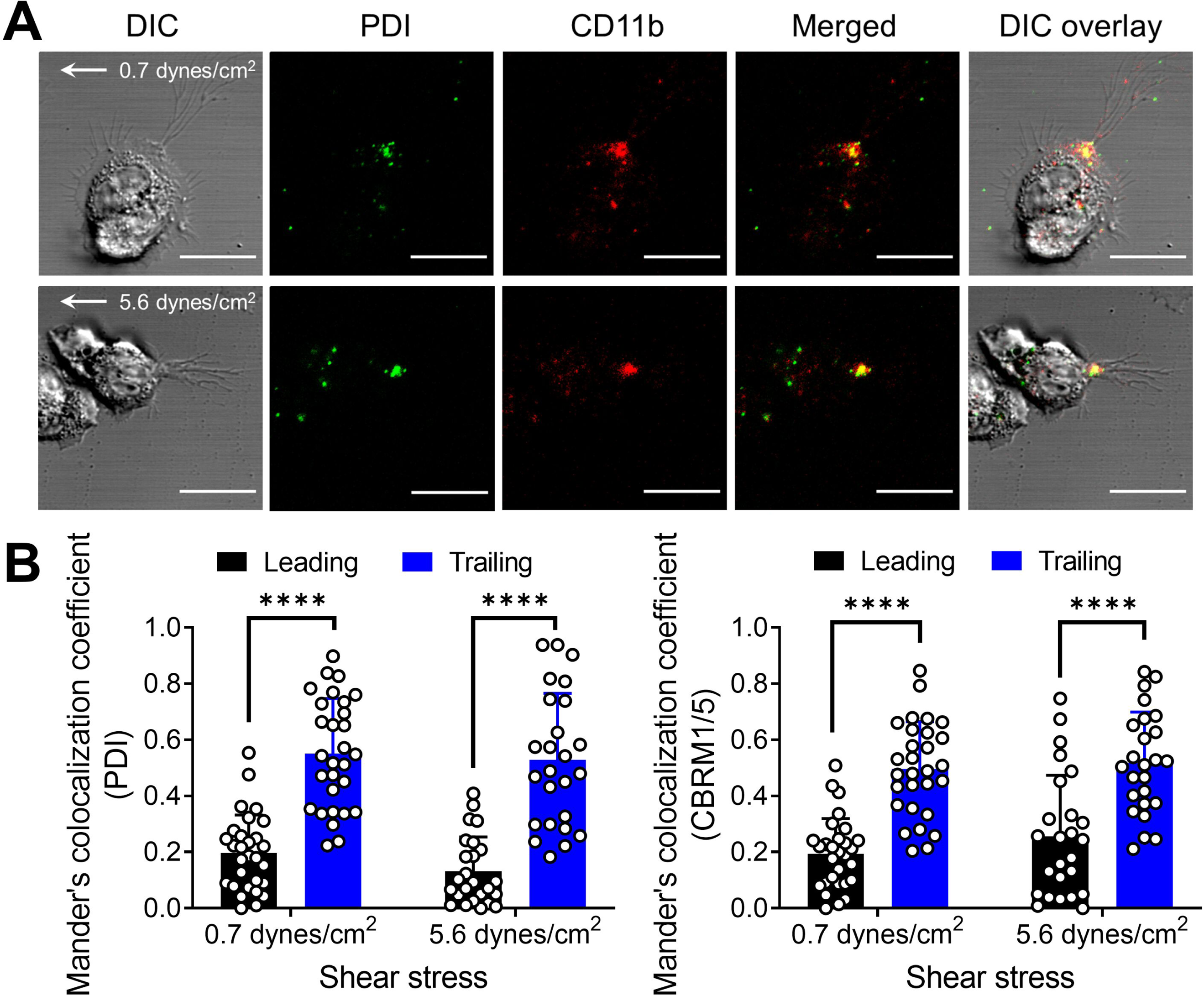
PDI colocalizes with active Mac-1 on trailing edge of neutrophils adhered to ICAM-1 under fluid shear. **(A)** Representative images of neutrophils adhered to ICAM-1 coated surface under fluid shear. Surface PDI was detected using anti-PDI antibody DL-11 and Alexa-Fluor 488-conjugated goat anti-rabbit IgG (green), and high affinity active Mac-1 was detected using an APC conjugated anti-CD11b antibody CBRM1/5 (red). After staining with antibodies, neutrophils were stimulated with 1 µM fMLF, perfused onto ICAM-1 coated microfluidic channels at 0.175 dynes/cm^2^ and left to adhere. Adhered neutrophils were subjected to fluid shear at 0.7 dynes/cm^2^ or 5.6 dynes/cm^2^ for 1 min before being imaged by confocal microscopy. Scale bar represents 10 µm. **(B)** Analysis of colocalization of surface PDI and Mac-1 at the leading or trailing edge of neutrophils. Leading and trailing edges of each neutrophil were defined by protrusion and trailing tail observed in DIC images. Manders’ colocalization coefficients for PDI and Mac-1 was determined from 29 cells at 0.7 dynes/cm^2^ and 25 cells at 5.6 dynes/cm^2^ from 3 independent experiments. Data shown is mean ± SD from independent experiments. ****P<0.0001 assessed by two-tailed, paired Wilcoxon test of the leading and trailing edges of the same cell.

Colocalization of PDI and Mac-1 in the trailing edge of crawling neutrophils suggests that PDI is manipulating disulfide bonds in Mac-1. This was measured using differential cysteine alkylation and mass spectrometry.

### PDI cleaves two disulfide bonds in the β2 integrin

Recombinant Mac-1 protein purified from human embryonic kidney cells was incubated with 10-fold molar excess of redox active PDI or redox inactive PDI (riPDI). Both catalytic cysteines in the *a* and *a’* domains were replaced with alanine in riPDI. Unpaired cysteines in Mac-1 were alkylated with the thiol alkylator 2-iodo-N-phenylacetamide (^12^C-IPA) followed by reduction of disulfide bonds by DTT and labeling of disulfide cysteines with a carbon-13 isotope of IPA (^13^C-IPA) (**Figure 2A**). The protein was digested by proteases and the peptides were analyzed by mass spectrometry. Forty-nine cysteine-containing peptides representing 24 of the 28 disulfide bonds in the β2 integrin subunit were detected and analyzed (**Figure 2B**, **Figure 2 – figure supplement 1** and **Supplementary File 1 Table S2**). Using existing crystal structures of extended αVβ3 (Xiong et al., 2009) and bent αXβ2 (Sen and Springer, 2016) and sequence alignment, a model of extended Mac-1 structure was constructed and the positions of the 28 disulfide bonds indicated (**Figure 2C**). The four disulfide bonds which were not resolved (C514-C537, C519-C535, C581-C590 and C593-C596) occur in the cysteine-rich EGF3 and EGF4 domains (**Figure 2C**). The redox state of the β2 disulfide bonds ranged from 90-98% oxidized in untreated control, which is in general agreement with the structure of mature β2 integrin where all disulfide bonds were found to be intact (Sen and Springer, 2016). Addition of redox active but not redox inactive PDI resulted in almost complete (>90%) and selective reduction of the C169-C176 and C224-C264 disulfide bonds in the βI domain (**Figure 2B**).

**Figure 2.**
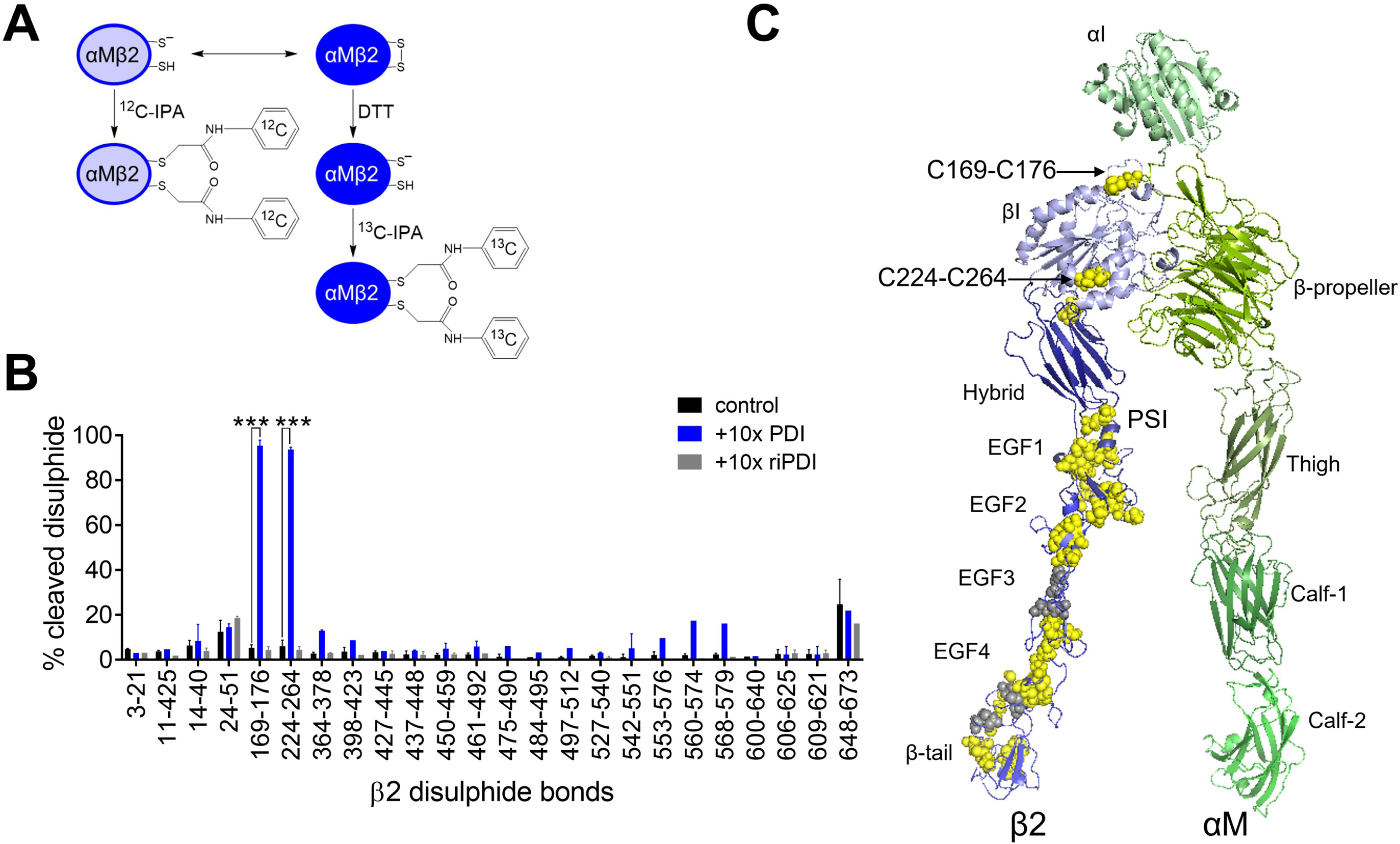
PDI selectively cleaves two βI-domain disulfide bonds in β2 integrin. **(A)** Differential cysteine alkylation and mass spectrometry method was employed to measure the redox state of disulfide bonds in recombinant β2 integrin. Unpaired cysteine thiols were labelled with ^12^C-IPA and the disulfide bonded cysteine thiols with ^13^C-IPA following reduction with DTT. 49 peptides encompassing 24 of the 28 β2 disulfide bonds were mapped. **(B)**. The redox state of 24 of the 28 β2 disulfide bonds was measured in the absence or presence of 10-fold molar excess of PDI or redox inactive PDI mutant (riPDI). Data shown is mean ± SEM of 2-4 peptides. ***P<0.001 assessed by unpaired, two-tailed Student’s t-test. **(C)** Model of Mac-1 in extended conformation. The β2 disulfide bonds quantified in panel B are shown as yellow spheres and disulfide bonds that were not mapped are shown as gray spheres. Both C169-C176 and C224-C264 disulfide bonds reside in the βI domain.

The β2 βI domain together with the β-propeller and I domain from the αM integrin subunit form the headpiece of the integrin. PDI cleavage of the βI domain disulfide bonds suggested that this vascular thiol isomerase might influence Mac-1 binding to ICAM-1.

### Ablation of the C224-C264 disulfide bond promotes Mac-1 de-adhesion from ICAM-1 under shear force

To study the effect of cleavage of the βI domain C169-C176 and C224-C264 disulfide bonds on binding of ligands to Mac-1, mammalian cells were transfected with wild-type or disulfide mutant integrins. One or both disulfide bonds were ablated by replacing the disulfide cysteines with serine, baby hamster kidney (BHK) cells stably transfected with wild-type αM and either wild-type β2 or mutant β2, and cells selected for comparable expression of the receptors (**Figure 3A**). Initially, Mac-1 binding to immobilized ICAM-1 in a static cell adhesion assay was assessed. Ablation of either of the two disulfide bonds had no effect on cell adhesion to ICAM-1 under static conditions (**Figure 3 – figure supplement 1**). As Mac-1 and ICAM-1 interact under shear forces in flowing blood to mediate neutrophil adhesion and crawling on the endothelium, we assessed Mac-1 binding to ICAM-1 under fluid shear. Two different states of PDI were employed in the assays; one where both active site cysteines were fully reduced (reduced PDI) and another where the active site cysteines were fully oxidized (oxidized PDI). BHK cells expressing wild-type Mac-1 were untreated or incubated with the different PDI forms before perfusing over ICAM-1-coated channels and left to adhere. Non-adherent cells were removed by perfusion of buffer at low shear force of 0.175 dynes/cm^2^. De-adhesion of bound cells was then triggered by doubling the shear force every minute until it reached 11.2 dynes/cm^2^. The number of cells remaining adhered at each shear force was measured (**Figure 3B**) and expressed as a percentage of the total adherent cells at 0.175 dynes/cm^2^. The area under the curve of the adherent cells as a function of shear force was calculated. Reduced but not oxidized PDI promoted de-adhesion of wild-type Mac-1 expressing cells from ICAM-1 (**Figure 3C**). The data was fit to a one phase exponential decay model by nonlinear regression to determine the decay constant (*K,* cm^2^ dynes^-1^) and shear force (*F_50_*, dynes/cm^2^) at which 50% of the cells were de-adhered from ICAM-1 (**Table 1**). The *F_50_* for wild-type Mac-1 expressing cells treated with reduced PDI is 0.7299 dynes/cm^2^, which is approximately half of the *F_50_* for cells treated with oxidized PDI or PBS vehicle. In other words, wild-type Mac-1 expressing cells require half the shear force to detach from ICAM-1 in the presence of reduced PDI compared to those in the presence of oxidized PDI. This result indicates that the disulfide-cleaving activity of PDI, but not its oxidizing activity, promotes Mac-1 de-adhesion from ICAM-1.

**Figure 3.**
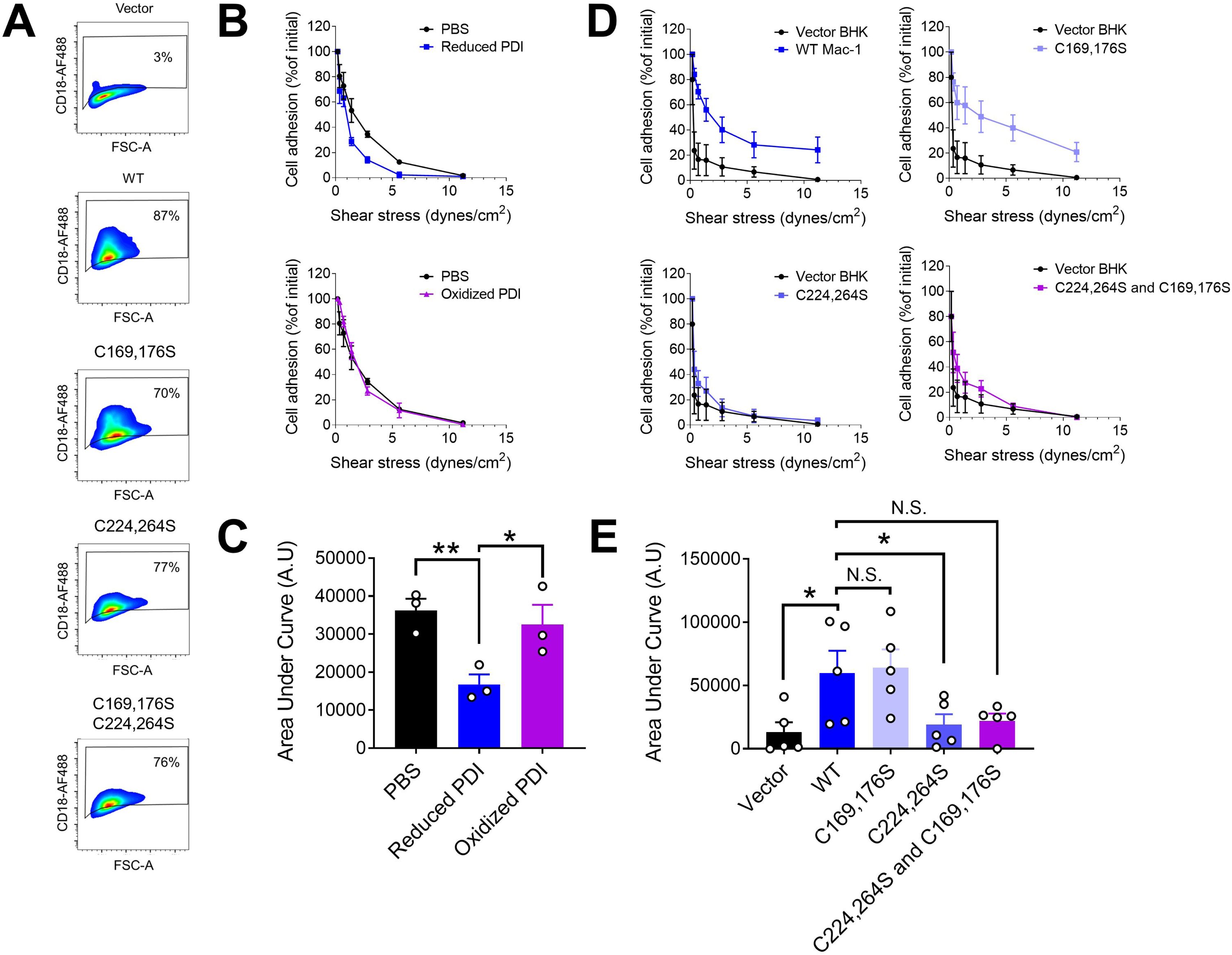
PDI cleavage of Mac-1 C224-C264 disulfide bond promote cell de-adhesion from ICAM-1 under shear force. **(A)** Detection of Mac-1 expression (WT or disulfide mutants) in BHK cells. cDNA constructs for *ITGAM* and *ITGB2* were co-transfected into BHK cells. Expression of wild-type (WT) Mac-1 or disulfide mutant were detected using Alexa Fluor 488 conjugated H52 antibody specific to β2 integrin by flow cytometry. Transfection with empty vector served as negative control. (**B**) De-adhesion assays of BHK cells expressing WT Mac-1 incubated without or with PDI to immobilized ICAM-1 under increasing shear force. Calcein-stained BHK cells incubated without or with 1µM reduced or oxidized PDI were perfused and allowed to adhere to ICAM-1 coated microfluidic channels for 15 min. Cells were subjected to shear force at each defined shear rate for 1 min (0.175, 0.35, 0.7, 1.4, 2.8, 5.6 and 11.2 dynes/cm^2^) to allow cell de-adhesion. Images were acquired and the number of adhered cells remained at each shear rate was quantified. **(C)** Area under each curve of de-adhesion from panel B. Data represent mean ± SEM of three biological replicates. *P<0.05; **P<0.01 by one-way ANOVA with Dunnett’s post-hoc multiple comparisons. **(D)** De-adhesion assays of BHK cells expressing WT Mac-1 or disulfide mutant to immobilized ICAM-1 under increasing shear force (0.175, 0.35, 0.7, 1.4, 2.8, 5.6 and 11.2 dynes/cm^2^). Images were acquired and the number of adhered cells remained at each shear rate was quantified. **(E)** Area under each curve of de-adhesion from panel D. Data represent mean ± SEM of three biological replicates. *P<0.05; N.S.=non-significant by one-way ANOVA with Dunnett’s post-hoc multiple comparisons.

**Table 1.** The decay constant (*K*) and shear force (*F_50_*) at which 50% of BHK cells expressing wild type or disulfide mutant Mac-1 were de-adhered from immobilized ICAM-1.

| | $K$ (cm<sup>2</sup>dynes<sup>-1</sup>) | $F_{50}$ (dynes/cm<sup>2</sup>) |
| --- | --- | --- |
| Vector alone | 8.979 | 0.07720 |
| WT Mac-1 + PBS | 0.4399 | 1.576 |
| WT Mac-1 + reduced PDI | 0.9496 | 0.7299 |
| WT Mac-1 + oxidized PDI | 0.4743 | 1.461 |
| WT Mac-1 | 0.5321 | 1.303 |
| C169,C176S | 0.6689 | 1.036 |
| C224,264S | 2.043 | 0.3392 |
| C224,264S and C169,C176S | 1.236 | 0.5606 |

To further define whether PDI promotes shear-dependent Mac-1 de-adhesion from ICAM-1 by cleavage of the two β2 disulfide bonds, we subjected BHK cells expressing wild-type Mac-1 or disulfide mutant Mac-1 to the same cell de-adhesion assays. Ablation of the C224-C264 disulfide bond but not the C169-C176 bond enhanced the shear-dependent de-adhesion of cells from ICAM-1 (**Figure 3D and E**). *F_50_* for cells expressing C224,264S Mac-1 is 0.3392 dynes/cm^2^, which is approximately one quarter of the *F_50_* for cells expressing wild-type Mac-1 (**Table 1**). This value is half of the *F_50_* for Mac-1 expressing cells treated with reduced PDI. The difference is possibly due to incomplete PDI cleavage of the C224-C264 disulfide bond in cell surface Mac-1. This finding indicates that PDI cleavage of the C224-C264 disulfide bond is important for Mac-1 dis-engagement from ICAM-1 under shear force.

The Mac-1 subpopulation reported to mediate ICAM-1 interaction in activated neutrophils has also been shown to bind fibrinogen (Diamond and Springer, 1993a). To characterize if PDI cleavage of β2 disulfide bonds promotes de-adhesion of Mac-1 from fibrinogen, we subjected BHK cells expressing wild-type or disulfide mutant Mac-1 to de-adhesion assays using fibrinogen-coated channels. Ablation of one or both β2 disulfide bonds had no significant effect on Mac-1 dis-engagement from fibrinogen when compared to wild-type Mac-1 (**Figure 3 – figure supplement 2**). This result indicates that PDI control of Mac-1 de-adhesion is selective for interaction with ICAM-1 under shear condition.

Integrin affinity for ligand is directly related to its conformations (Chen et al., 2010; Diamond and Springer, 1993b). Mac-1 transitions from bent closed conformation to open extended conformations that correlate with transition from low- to high-affinity state for ligand engagement. Intermediate extended closed conformations have also been observed in leukocyte integrins (Fan et al., 2019) (**Figure 4A**).

**Figure 4.**
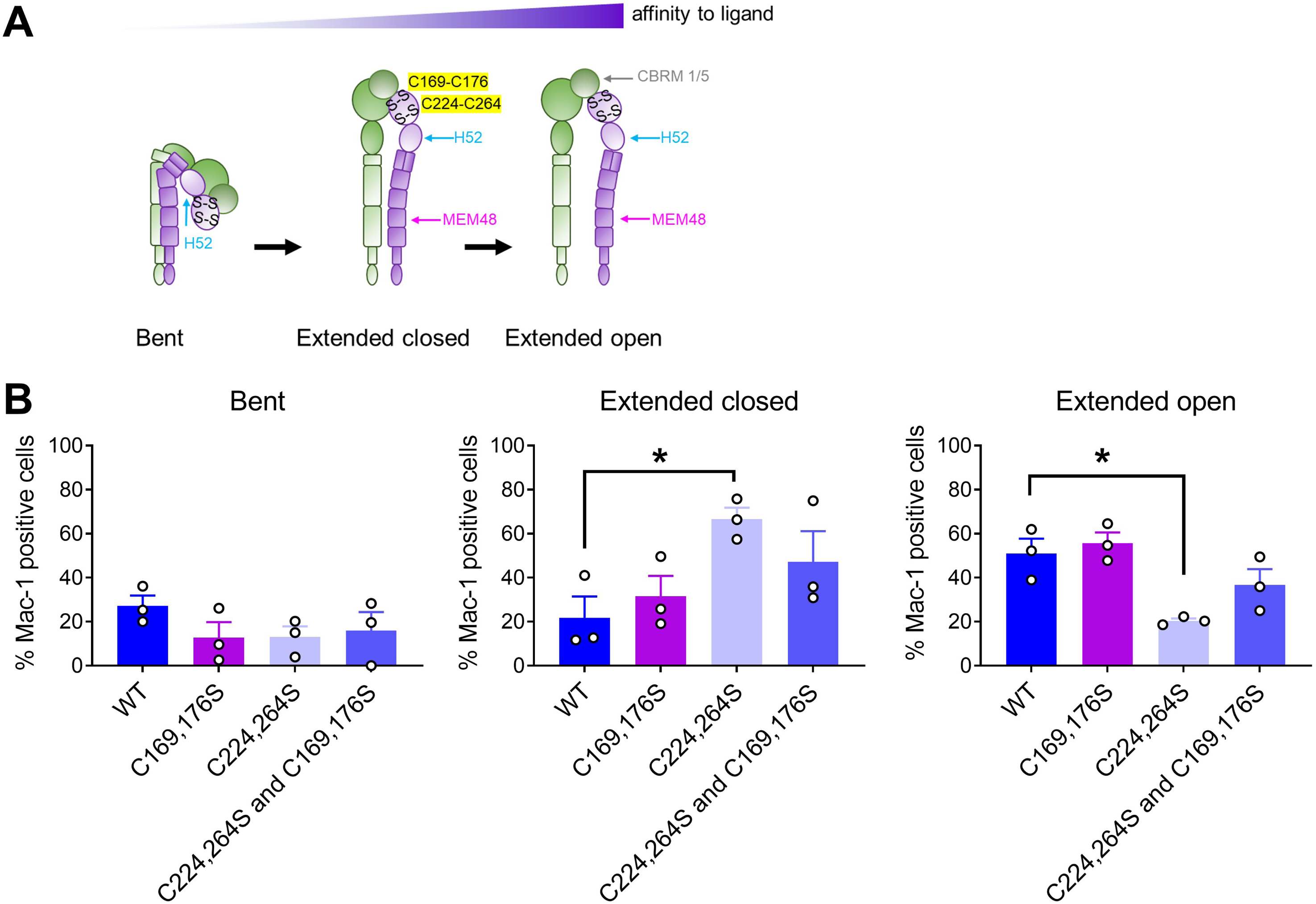
Cleavage of the C224-C264 disulfide bond favors Mac-1 extended closed conformation. **(A)** Schematic representation of the Mac-1 conformation states and the location of epitopes recognized by antibodies H52 (conformation independent), MEM48 (extended state) and CBRM1/5 (extended open state). The two βI-domain disulfide bonds (S-S) are shown. **(B)** Determining the conformations of Mac-1 (WT or disulfide mutants) expressed in BHK cells. The total number of cells expressing Mac-1 was determined using H52 antibody. The proportion of cells expressing extended (both closed and open) Mac-1 was measured using MEM48 antibody, while cells expressing extended open state of Mac-1 was determined using CBRM1/5. Proportion of cells expressing extended closed conformation was calculated by subtracting the number of CBRM1/5 positive cells from the MEM48 positive cells. Proportion of cells expressing bent conformation was calculated by subtracting the MEM48 positive cells from the H52 positive cells. Data shown is mean ± SEM of three independent experiments. *P<0.05 assessed by one-way ANOVA, with Dunnett’s post-hoc multiple comparisons

### Ablation of the C224-C264 disulfide bond favors a lower affinity state of Mac-1

To investigate how ablation of the β2 βI domain C224-C264 disulfide bond influences Mac-1 conformations and affinity states, we employed conformation reporting antibodies and flow cytometry to probe the distribution of conformations of wild-type and disulfide mutant Mac-1 expressed on BHK cells.

Total Mac-1 expression on BHK cells was determined using the H52 monoclonal antibody that recognizes an epitope in the hybrid domain (residues 386-400) of the β2 subunit (Al-Shamkhani and Law, 1998) and is accessible in all Mac-1 conformations (**Figure 4 – figure supplement 1**). The monoclonal antibody, MEM48, recognizes the EGF3 domain (residues 534-543) of β2 integrin that is only exposed when Mac-1 is extended (Sen and Springer, 2016). The monoclonal antibody, CBRM1/5, recognizes an epitope in the αM I domain (P147, H148, R151, K200, T203, L206) that becomes exposed in the fully extended open conformation (Oxvig et al., 1999). The ratio of CD11b+ to CD18+ cells for cells expressing wild-type, C224,264S and C224,264S and C169,176S mutant Mac-1 ranges from 0.8 to 1.1 indicating comparable expression of the receptor forms (**Supplementary File 1 Table S3**). For the C169,176S Mac-1 mutant, the ratio is 1.5-1.7 and the number of H52+ cells were normalized according to the CD11b+ to CD18+ ratio. There was no significant difference in the distribution of bent (10-30%), extended and closed (20-40%), and extended open (50-60%) conformations in cells expressing wild-type, C169,176S mutant, and C224,264S and C169,176S double mutant Mac-1 (**Figure 4B**). In contrast, there was a shift of conformations from extended open (20%) to predominately extended closed (70%) in cells expressing C224,264S mutant Mac-1. In other words, the ablation of the C224-C264 bond altered Mac-1 conformation to favor a lower affinity state for ligand binding.

To elucidate how the β2 βI C224-C264 bond could influence ligand affinity, we conducted MD simulations of the effect of C169-C176 and C224-C264 redox state on the conformational dynamics of the βI domain in complex with the β-propeller (**Figure 5A**).

**Figure 5.**
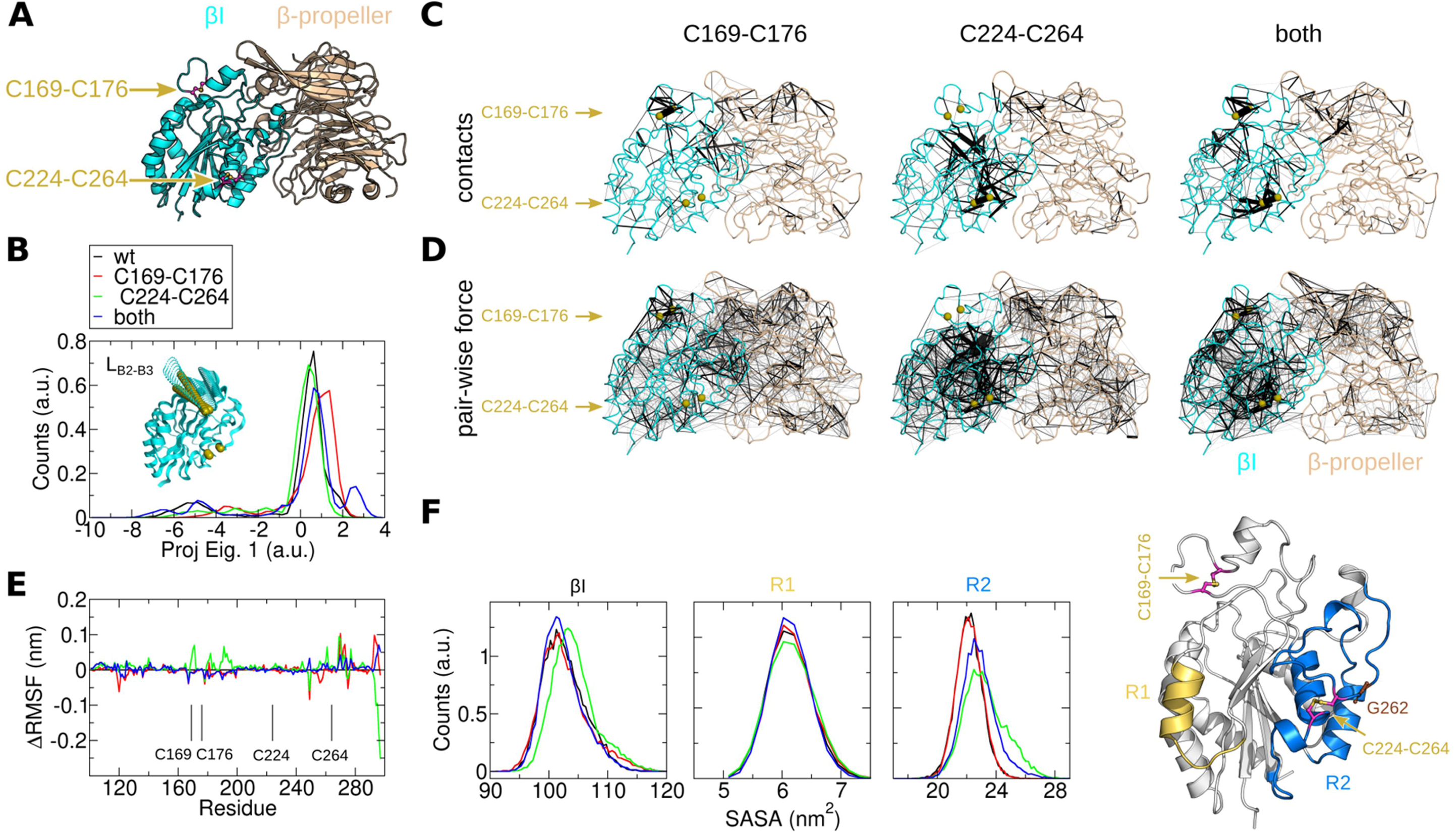
Cleavage of the C224-C264 disulfide bond perturbs inter-residue contact and mechanical in the βI domain of β2 integrin. **(A)** Structure of the complex is shown (β-propeller in wheat and βI domain in cyan). Position of the C169-C176 and C224-C264 disulfide bonds is indicated. Atomic positions for simulations were taken from the X-ray structure of the Leukocyte integrin ɑLβ2 (PDB identifier 5E6U^1^). LFA-1 (ɑLβ2) and Mac-1 (ɑMβ2) have identical βI domains and highly similar β-propeller domains (73% sequence similarity, 41% residues being identical). **(B)** Protein conformational changes of βI were monitored by principal component analysis (PCA). The main PCA eigenvector (Eig. 1) was related to conformational transitions of the loop which connects strands B2 and B3 (inset). MD trajectories were projected onto this eigenvector and the resulting histograms of the projections are shown for each simulated condition. **(C)** Change in the pair-wise contact probability ΔC_ij_ is mapped on the 3D structure of the complex, by lines connecting the C-alpha atoms of residues i and j. The thickness of the line is proportional to the absolute value of ΔC_ij_, ranging from 0 to 0.4. Changes larger than 0.4 have the same (maximum) line thickness. Changes are presented for the system, X, with the C169-C176 bond (left), the C224-C264 bond (middle) bond, or both bonds (right) reduced, with respect to the wild-type situation (wt) in which both disulfide bonds are formed: ΔC_ij_=C_ij_(X)-C_ij_(wt). **(D)** Change in pair-wise force ΔF_ij_ with the same format as in C. ΔF_ij_, varies from 0 to 100 pN, and changes larger than the latter have the same (maximum) line thickness. In C and D, pairs with normalized difference z>0.5 are shown. See definition of z in the methods and the effect of the z value in **Supplementary Fig. 6. (E)** Change in root mean square fluctuation (RMSF) versus amino acid sequence of βI is displayed: ΔRMSF=RMSF(X)-RMSF(wt), with X corresponding to the four simulated systems (same color coding as in B). Positions along the sequence of the involved cysteines are highlighted with the vertical lines. **(F)** Histograms of the solvent accessible surface area of βI (left panel) and some regions of it (middle and right panels) for the four studied systems are shown (same color as in B). Region R1 corresponds to the amino acids 188–200 (yellow) and region R2 to amino acids 214–234 and 256–293 (blue). Both regions are highlighted in the cartoon representation of βI at the right side.

### Cleavage of the C224-C264 bond perturbs inter-residue contact and mechanical stress in the βI domain of the β2-integrin

As the structure of Mac-1 has not yet been determined, we took initial atomic positions from the X-ray structure of the highly close homolog, LFA-1 (ɑLβ2, CD11a/CD18) (Sen and Springer, 2016). LFA-1 has an identical βI domain and highly similar β-propeller to Mac-1 (**Figure 5A**). The dynamics of the complex was monitored in multiple molecular dynamics simulation replicas and for different redox states of the C169-C176 and C224-C264 disulfide bonds. During the simulations, the complex was found to be very stable, with a backbone root mean square deviation from the initial positions smaller than 0.45 nm. Inside the βI domain, the loop connecting the strands B2 and B3 (L_B2-B3_) displayed the largest conformational variations, although the redox state of the bonds of interest did not favor any preferential position of this loop (**Figure 5B**). We also analyzed the change in residue-residue contacts induced by reduction of either disulfide bond (**Figure 5C** and **Figure 5 – figure supplement 1**). Reduction of C224-C264 altered the contact probability of more residue pairs than reduction of C169-C176 did. Reduction of C169-C176 bond resulted in perturbations nearby the disulfide, while perturbations extended to other regions of βI when the C224-C264 bond was reduced. Reduction of both bonds at the same time showed a different perturbation pattern, with changes in the contact probability close to both affected bonds but not in the region between them. In addition, we examined how disulfide bond reduction altered the internal mechanical stress of the protein. To this end, we computed changes in the residue-residue pair-wise forces (**Figure 5D** and **Figure 5 – figure supplement 1**). Consistent with the residue-residue contacts, the pair-wise force pattern changed much more drastically upon reduction of C224-C264 than reduction of C169-C176, or after reduction of both bonds. Moderate changes in the internal mechanical stress propagate beyond the βI domain, even reaching the β-propeller, while more pronounced differences occurred mainly in the βI domain (compare the situation for moderate changes, z>0.5, with that for large changes, z>0.75 and z>1.0, in **Figure 5 – figure supplement 1**).

The question arises how this allosteric effect originating from C224-C264 reduction impacts the complex structurally. Calculation of the root mean square fluctuation (RMSF) also displayed changes in the dynamics of several residues distant to C224-C264, even close to C169-C176 (**Figure 5E**). In addition, the solvent accessible surface area (SASA) of βI was found to shift towards larger areas when the C224-C264 bond was reduced, but not in the other situations (**Figure 5F**). This increment in SASA is attributed to a more exposed surface area of the region R2 near C224-C264 rather than the distant region R1 for which changed connectivity and stress was also observed (compare regions R2 and R1 in **Figure 5F**). In summary, our MD simulations demonstrate that reduction of the C224-C246 bond, and to a minor extent the reduction of C169-C176 or both, allosterically alters the internal connectivity and mechanical stress and modulates the surface area of βI.

To demonstrate how PDI and force are essential to regulate neutrophils de-adhesion from ICAM-1, we measured neutrophil crawling as a function of cell adhesion and de-adhesion events under fluid shear.

### PDI promotes neutrophil crawling in the direction of flow

Neutrophils express two integrins, LFA-1 and Mac-1, that both interact with ICAM-1. Genetic knockout of PDI from neutrophils, however, only impairs Mac-1 function but not LFA-1 (Hahm et al., 2013). It was recently reported that Mac-1 is essential for neutrophil migration in the direction of flow while LFA-1 mediates movement against flow when Mac-1 is inhibited by function blocking antibodies (Buffone et al., 2019). To determine how PDI cleavage of Mac-1 disulfide bond influences neutrophil migration, neutrophils were treated with control oxidized or active reduced PDI, stimulated with fMLF and perfused over an ICAM-1 coated surface. Adhered neutrophils were then subjected to 0.7 or 5.6 dynes/cm^2^ fluid shear and cell tracks measured (**Figure 6A**). Displacement of neutrophils in X and Y directions from their initial position was determined (**Figure 6 – figure supplement 1**; **Supplementary Videos 1-4**) and expressed as migration index, which is defined as the ratio of the difference between the initial and final X- or Y-displacement over the total distance travelled by a neutrophil. Positive values for migration index in the X-direction indicate cell displacement with flow while negative values indicate cell displacement against flow. Zero indicates no preferred direction. When subjected to 0.7 dynes/cm^2^ fluid shear, neutrophils treated with control oxidized or active reduced PDI show no significant difference in their migration in the X-direction (**Figure 6B**). In other words, PDI had no effect on neutrophil migration with or against flow at 0.7 dynes/cm^2^. Percentage of neutrophils migrating in the direction of flow at 0.7 dynes/cm^2^ was 32% for cells treated with oxidized PDI and 42% for cells treated with reduced PDI (**Figure 6 – figure supplement 2; Supplementary File 1 Table S4**). In contrast, when subjected to 5.6 dynes/cm^2^ fluid shear, there was a significant increase of neutrophils migrating in the direction of flow when treated with reduced PDI when compared to neutrophils treated with control oxidized PDI (**Figure 6B**). Percentage of neutrophils migrating in the direction of flow at 5.6 dynes/cm^2^ was 42% for cells treated with oxidized PDI and increased to 70% for cells treated with reduced PDI (**Figure 6 – figure supplement 2; Supplementary File 1 Table S4**). Displacement of neutrophils in the Y-direction which is perpendicular to the direction of flow was also determined (**Figure 6C**). Neutrophils treated with oxidized or reduced PDI had no significant difference in their migration in the Y-direction, indicating that PDI has no effect on neutrophil movement in the direction perpendicular to fluid shear.

**Figure 6.**
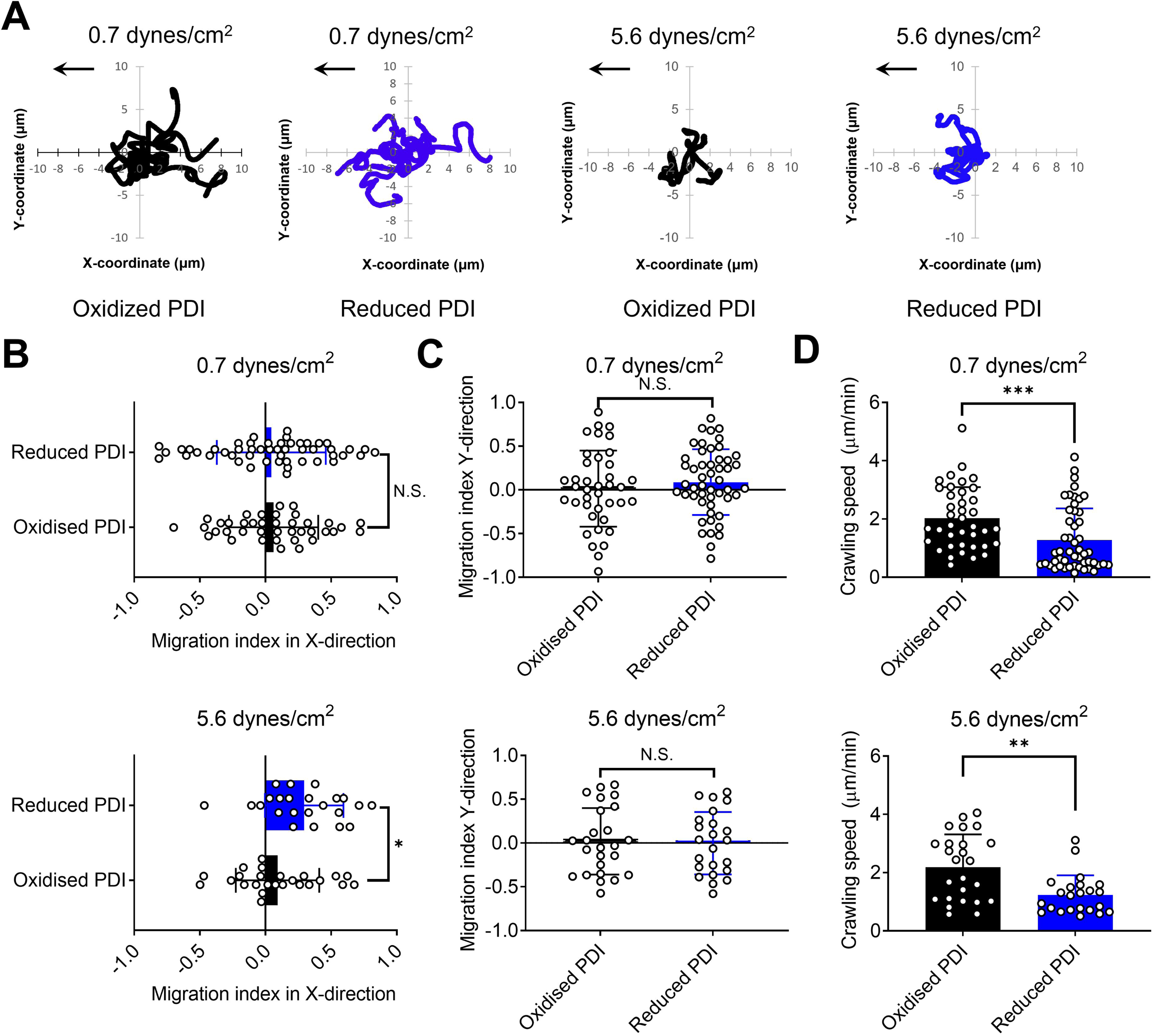
Reduced PDI but not control oxidized PDI promotes neutrophil motility in the direction of flow at high shear stress. **(A)** Cell tracks of neutrophils migrating in 0.7 dynes/cm^2^ or 5.6 dynes/cm^2^ fluid shear were measured to determine distance and direction of travel during the experiment. Each plot represents a track from an individual neutrophil. Arrow indicates the direction of shear force. Directional migration of neutrophils under fluid shear is expressed as migration index in **(B)** X- or **(C)** Y-direction. Neutrophils were treated with control oxidized or reduced PDI, stimulated with fMLF and perfused over ICAM-1-coated microfluidic chips. Adhered neutrophils were subjected to 0.7 or 5.6 dynes/cm^2^ fluid shear and their X- and Y-displacement and total distance traveled was measured for calculation of migration index for each cell. A negative migration index in the X-direction indicates cell migration against the flow whereas a positive migration index in the X-direction indicates cell migration with the flow. **(D)** Crawling speed of neutrophils at 0.7 or 5.6 dynes/cm^2^ shear force was calculated by determining the total distance traveled over the total time of migration. Data was mean ± SD of three independent experiments. A total of 40 and 25 cells treated with control oxidized PDI and 50 and 23 cells treated with reduced PDI was analyzed for shear force at 0.7 dynes/cm^2^ and 5.6 dynes/cm^2^ fluid shear, respectively. *P<0.05; **P<0.01; ***P<0.001; N.S.=non-significant assessed by unpaired, Mann-Whitney test.

Crawling speed of neutrophils treated with control oxidized or reduced PDI was also determined by measuring the total distance traveled by each neutrophil and dividing it by the total time of migration. Neutrophils treated with reduced PDI exhibited significantly slower crawling speeds at 0.7 and 5.6 dynes/cm^2^ fluid shear compared to neutrophils treated with control oxidized PDI (**Figure 6D**), and the speeds were comparable at both shear force (**Supplementary File 1 Table S5**). This result indicates that PDI-mediated decrease in crawling speed of neutrophils treated with is independent of shear force.

Together, our data indicates that reduced PDI but not control oxidized PDI slows down neutrophil crawling under shear force and promotes migration in the direction of flow.

## Discussion

We describe here a mechano-redox event controlling the function of Mac-1 on the trailing edge of neutrophils. PDI in the presence of fluid shear from 0.17-11 dynes/cm^2^ selectively regulates Mac-1 de-adhesion from endothelial ICAM-1 by cleaving the C224-C264 allosteric disulfide bond in the β2 βI domain (**Figure 7A**). Cleavage of this bond induces mechanical stress in the βI domain and allosterically perturbs residue contacts between the βI and β-propeller domains. We suggest that this conformational change in Mac-1 results in suboptimal binding to ICAM-1 that leads to detachment of ICAM-1 in the fluid shear encountered in the circulation. As a consequence of Mac-1 de-adhesion at the trailing edge of the cell, PDI promotes neutrophil migration in the direction of flow.

**Figure 7.**
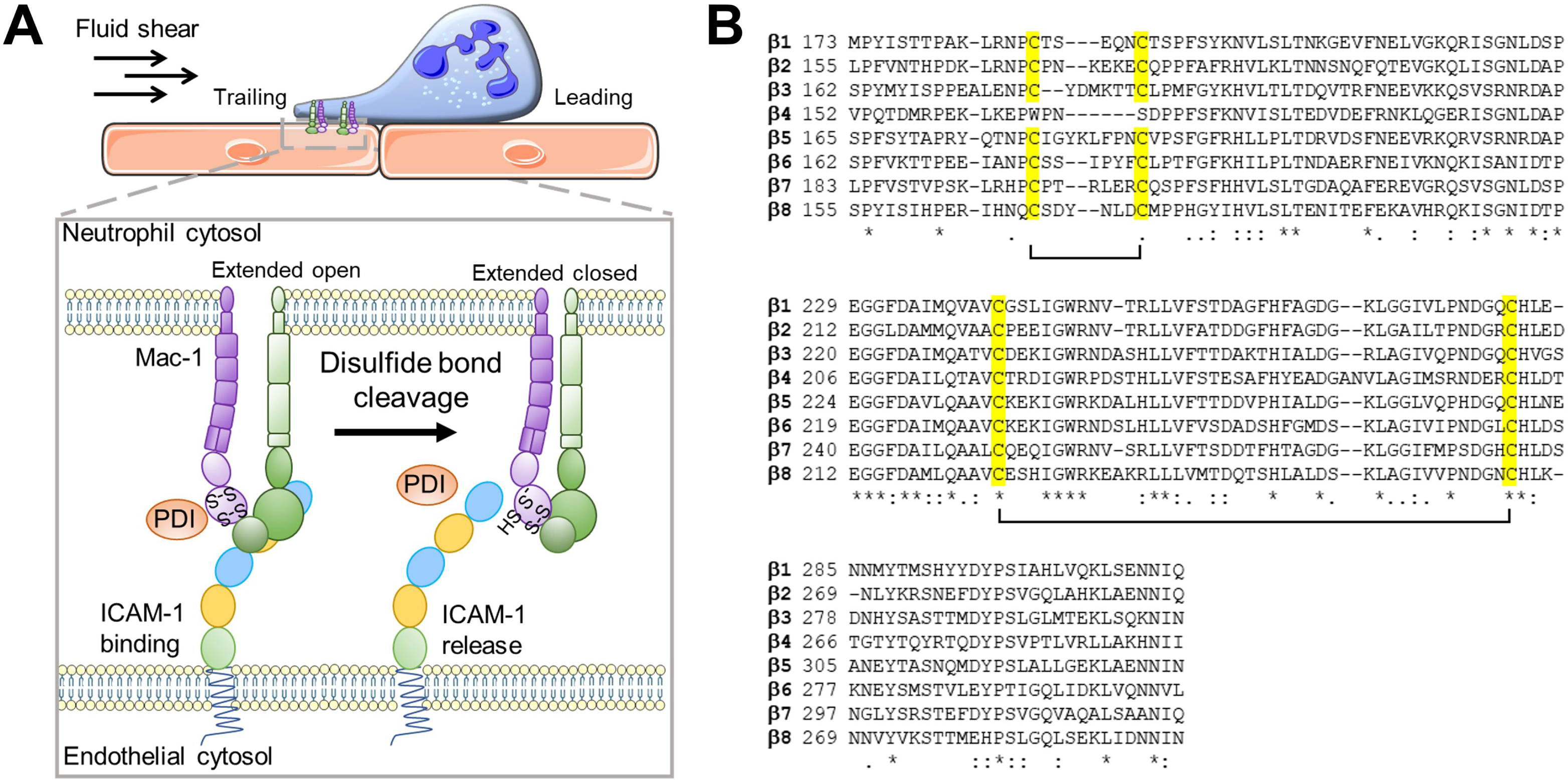
Schematic representation of mechano-redox control of Mac-1 de-adhesion from ICAM-1 by PDI cleavage of an allosteric disulfide bond at the trailing edge of neutrophils. In its extended and open conformation, Mac-1 mediates high affinity adhesion of neutrophils to endothelial cells via binding to ICAM-1 under shear stress. PDI colocalizes with high affinity Mac-1 at the trailing edge of neutrophils and regulates neutrophil adhesion to endothelial cells by cleaving the βI-domain disulfide bond. PDI cleavage of this disulfide induces internal mechanical stress in the βI domain leading to Mac-1 switch from an extended open to an extended closed conformation with lower affinity for ICAM-1. This results in de-adhesion at the trailing edge of neutrophils to promote migration on endothelium in the direction of flow. **(B)** Alignment of the βI domain from human β integrins. Protein sequences of integrins β1-8 were obtained from Uniprot and aligned using Clustal Omega (Sievers et al., 2011). Conserved cysteines are highlighted in yellow and cysteine pairings for disulfide bonds are indicated by brackets.

Leukocyte integrins, LFA-1 and Mac-1, are major integrins that interact with endothelial ICAM-1 to mediate cell motility. In T-cells and hemopoietic stem cells, the predominant expression of LFA-1 enables these cells to migrate on the endothelium against the direction of flow (Buffone et al., 2018). However, in neutrophils which express both LFA-1 and Mac-1, cells migrate only in the direction of flow. Mac-1 expression is the key determinant for neutrophil directional migration since inhibition of Mac-1 by function-blocking antibodies results in neutrophil migration against flow (Buffone et al., 2019). How Mac-1 mediates neutrophil movement with flow has been elusive. In a crawling neutrophil, LFA-1 localizes in the trailing edge while Mac-1 localizes in both the trailing edge and in the middle of the cell (Hyun et al., 2019). By measuring separation of the cytoplasmic tails of α and β subunits using fluorescence resonance energy transfer, LFA-1 is found to adopt a high affinity state when bound to ICAM-1 in a moving neutrophil whereas Mac-1 adopts a lower affinity state (Hyun et al., 2019). The results described herein indicate that the lower affinity of Mac-1 for ICAM-1 in a moving neutrophil can be attributed to PDI cleavage of the Mac-1 C224-C264 disulfide bond that leads to shift in Mac-1 conformation from extended open to extended closed state. Our colocalization data indicates that PDI selectively targets Mac-1 clusters at the trailing edge of neutrophils. It has been suggested that the uropod senses the direction of flow via the trailing edge, which is less adherent and therefore more susceptible to shear force (Valignat et al., 2014). Our findings provide a molecular mechanism for neutrophil detachment at the trailing edge that enables sensing of directional flow. Shear-dependent PDI cleavage of the Mac-1 C224-C264 allosteric disulfide bond spatially regulates Mac-1 affinity at the trailing edge to drive neutrophil movement in the direction of flow.

We also observed PDI-dependent reduction in the crawling speed of neutrophils that was independent of shear force. Disruption of binding of Mac-1 to ICAM-1 by Mac-1 blocking antibodies reduces the crawling speed of neutrophils *in vitro* and *in vivo* (Volmering et al., 2016) (Li et al., 2018), which is consistent with our observations. That is, PDI cleavage of the Mac-1 C224-C264 disulfide bond phenocopies Mac-1 function blocking antibodies in disrupting Mac-1 adhesion to ICAM-1. It has been reported that proteases released from the uropod mediate neutrophil detachment under static conditions by degrading surface Mac-1 (Singh et al., 2012). In our studies, PDI has no effect on Mac-1 adhesion to ICAM-1 in static conditions, which suggests there may be separate mechanisms regulating neutrophil migration under static versus shear conditions.

We previously reported that platelet surface thiol isomerase ERp5 cleaves the βI domain C177-C184 disulfide bond in the β3 subunit of platelet αIIbβ3 and this cleavage changed the positions of residues critical for metal ion coordination resulting in release of fibrinogen. The β2 subunit C169-C176 disulfide bond in Mac-1 is homologous to the β3 subunit C177-C184 disulfide bond (**Figure 7B**) and is close to the metal ion binding sites and αM I (or αI) domain involved in ICAM-1 binding. We, therefore, anticipated that the redox state of this bond would be a critical determinant of binding of Mac-1 to ICAM-1. Although our mass spectrometry analysis showed that PDI cleaves both βI-domain disulfide bonds equally well, our functional data supports that only the C224-C264 disulfide bond controls Mac-1 affinity for ICAM-1 in fluid shear. This finding is surprising since the C224-C264 disulfide bond is distant from known epitopes in the αI, β-propeller and βI domain essential for ICAM-1 interaction (Bajt et al., 1995; Chen et al., 2010; Diamond et al., 1991; Oxvig et al., 1999; Sen and Springer, 2016; Sen et al., 2013; Shimaoka et al., 2003; Yang et al., 2004). Our data using conformation reporting antibodies revealed that the redox state of the C224-C264 disulfide bond controls exposure of critical residues (P147, H148, R151, K200, T203, L206) in the αI domain required to form a fully open ligand binding pocket for high affinity ICAM-1 binding. This finding indicates that the C224-C264 disulfide bond influences the αI domain in an allosteric manner. Indeed, our MD simulations showed that cleavage of the C224-C264 disulfide bond perturbs contacts of both neighboring and distal residues thus supporting an allosteric mechanism of control.

Unlike LFA-1 which readily interacts with ICAM-1 regardless of its conformation state, Mac-1 only binds to ICAM-1 in its fully open extended conformation (Li et al., 2013). This suggests that αI domain alone may be insufficient for high affinity binding of Mac-1 to ICAM-1 but requires contact from the βI domain. Mutations identified in the deleterious genetic disease, Leukocyte Adhesion Deficiency Type I (LAD-1), support this hypothesis. Mis-sense mutations for a number of residues in R2 region can lead to LAD-1 (van de Vijver et al., 2012). One of the most common LAD-1 alleles is G284S (or G262S in the mature β2 integrin) that is two residues from C264. G284 (or G262) is precisely at the region that displayed the largest change in area exposed upon reduction of the C224-C264 disulfide bond. Expression of Mac-1 containing the G284S mutation in CHO cells resulted in reduced Mac-1 expression while a G284R (or G262R) mutation was associated with impaired ICAM-1 binding (Mathew et al., 2000; Uzel et al., 2008). These findings support our conclusion that residues influenced by the redox state of the C224-C264 disulfide bond are important for ICAM-1 binding.

Mechano-redox regulation of Mac-1 by PDI controls ICAM-1 but not that of fibrinogen binding. Among the integrin family, Mac-1 is considered the most promiscuous that can bind to over 30 extracellular ligands (Hyun et al., 2009). Distinct epitopes in Mac-1 have been identified to be important for binding to specific ligands (Diamond et al., 1993). For example. a motif in the αI domain M7 (E162-L170) specifically interacts with the inflammatory ligand CD40L. Inhibitory anti-M7 antibody blocks Mac-1 binding to CD40L but has no effect on binding to ICAM-1 or fibrinogen (Wolf et al., 2018). On the other hand, the αI domain epitopes for ICAM-1 and fibrinogen binding are overlapping as demonstrated by blocking of Mac-1 adhesion to both ICAM-1 and fibrinogen by anti-αM CBRM1/5 antibody (Diamond and Springer, 1993a). PDI, therefore, selectively controls Mac-1 promiscuity. Notably, the βI-domain C169-C176 and C224-C264 disulfide bonds are conserved in 7 of the 8 β integrins (**Figure 7B**) suggesting that other β integrins might be also subject to mechano-redox regulation.

In conclusion, we have identified a mechano-redox mechanism that selectively controls Mac-1 binding to ICAM-1 in fluid shear conditions. This mechanism allows the trailing edge of neutrophils to detach from ICAM-1 and enables movement in the direction of flow. Importantly, this informs studies on the development and optimization of PDI inhibitors as therapeutic agents to attenuate neutrophil migration to subdue inflammation.

## Materials and Methods

### Neutrophil Isolation

All procedures involving human whole-blood were collected from healthy human volunteers in accordance with the Human Research Ethics Committee of the University of Sydney (2014/244) and the declaration of Helsinki. Human whole-blood was collected from healthy human volunteers into plastic syringes containing Clexane at 20 U/mL (Sanofi). Neutrophils were isolated from human whole-blood by Histopaque density gradient centrifugation. A 48 mL density gradient was created by layering Histopaque-1077 on top of Histopaque-1191 (Sigma-Aldrich), followed by a layer of whole blood at a volumetric ratio of 2:1:1 in a 50 mL conical centrifuge tube at 25°C. The tube was centrifuged at 600 G for 20 min with no brake, and the pink neutrophil buffy coat that formed above the red blood cell layer was collected, diluted in Hank’s Buffered Salt Solution (HBSS, 1 mM CaCl_2_, 1 mM MgCl_2_, 5.4 mM KCl, 0.44 mM KH_2_PO_4_, 136.9 mM NaCl, 0.34 mM Na_2_HPO_4_, 5.5 mM D-glucose, 0.5% (w/v) BSA, pH 7.2), and centrifuged at 600 G for 5 min. The cell pellet was then resuspended in HBSS and cleared of red blood cells by lysis in ice cold 0.2% (w/v) NaCl for 20 secs, neutralized with an equivalent volume of 1.6% (w/v) NaCl and centrifuged at 250 g, 4°C for 6 min. This process was repeated up to 2 times until all red blood cells were cleared. Neutrophils were stored at 4°C, brought up to 25°C prior to use in assays, and used within 4 h of blood collection.

### Colocalization of Mac-1 and PDI on neutrophils

For colocalization of Mac-1 and PDI under static conditions, wells of an 8-well microslide were coated with 10 μg/mL of ICAM-1/Fc (R&D systems) for 2 h at room temperature, washed with PBS and blocked with 0.5% (w/v) BSA for 30 min. After a final wash, neutrophils (1×10^6^ cells/mL) were added to each well in the presence of APC conjugated anti-CD11b antibody CBRM1/5 (BioLegend) at 1 μg/mL, anti-PDI antibody DL11 (Sigma-Aldrich) at 2 μg/mL, and Alexa Fluor 488 conjugated goat anti-rabbit IgG (Thermofisher) at 2 μg/mL. Neutrophils were added to the wells without or with 10 μM of fMLF and left to adhere for 30 min at 37°C. After incubation, neutrophils were fixed with 4% (w/v) PFA for 1 h, washed with PBS, and covered with ProLong gold Antifade reagent (Thermofisher) according to manufacturer’s instructions before imaged on a Zeiss LSM880 confocal microscope with a 63x oil objective (NA 1.4).

For colocalization of Mac-1 and PDI under fluid shear, neutrophils were incubated with 1 μg/mL of an APC-conjugated anti-CD11b CBRM1/5 and 2 μg/mL of the rabbit anti-PDI DL-11 for 1 h on ice. Neutrophils were then washed and stained with 0.5 μg/mL of an Alexa Fluor 488-conjugated goat anti-rabbit IgG for 1 h on ice. After a final wash with Hank’s buffered saline solution, neutrophils were primed with 1 μM of fMLF, perfused through microfluidic devices coated with 10 μg/mL of ICAM-1/Fc, and left to settle for 5 min. Neutrophils were then exposed to 0.7 dynes/cm^2^ (100 s^-1^) or 5.6 dynes/cm^2^ (800 s^-1^) of shear by perfusing Hank’s buffered saline solution containing 5 μM of fMLF and imaged using a 63x oil objective (1.4 NA) on a Zeiss LSM880 confocal microscope.

Colocalization was analyzed using ImageJ. Regions of interest were drawn around the trailing and leading edge of crawling neutrophils, and the Coloc2 plugin was used to calculate the Pearson’s correlation coefficient and Mander’s coefficients.

### Redox state of disulfide bonds in β_2_ integrin

Recombinant redox active and redox inactive PDI were produced from *E.coli* as described (Passam et al., 2018). PDI was reduced by incubating with 10 mM DTT for 30 min at 25°C prior use. Reduced PDI was then desalted to remove DTT using 7K MWCO Zebaspin columns (Thermofisher). 2 µg of recombinant Mac-1 integrin (R&D systems) was incubated with reduced PDI or enzymatically inactive PDI at 10 µM for 30 min at 25°C. Reduced cysteines were then alkylated with ^12^C-IPA (Cambridge Isotopes) at 4 mM, 10% DMSO, for 1 h at 25°C. The αM and β2 subunits of recombinant Mac-1 integrin were resolved on 4-20% polyacrylamide (BioRad), gradient gels by SDS-PAGE and stained with Coomassie brilliant blue R250. The bands corresponding to the αM and β2 subunits were excised from the gels, destained, dried, and incubated with DTT at 40 mM for 30 min at 56°C and washed. The fully reduced protein was then alkylated with ^13^C-IPA at 4 mM in 10% DMSO, for 1 h at 25°C. The gel slices were washed and deglycosylated with 5 units of PNGase F (Sigma-Aldrich) overnight at 37°C. The proteins were digested with 12.5 ng/µL of chymotrypsin in 25 mM NH_4_HCO_3_, 10 mM CaCl_2_ for 4 h at 37°C, followed by digestion with 12.5 ng/mL trypsin in 25 mM NH_4_HCO_3_ overnight at 25°C. Peptides were eluted twice from the gel pieces with formic acid 5% v/v in acetonitrile 50% v/v. Liquid chromatography, mass spectrometry and data analysis were performed as previously described (Chiu, 2019). Briefly, peptides were analyzed on a Thermo Fisher Scientific Ultimate 3000. Two hundred ng of peptides was injected and resolved on a 35 cm × 75 μm C18 reverse phase analytical column with integrated emitter using a 2-35% acetonitrile over 20 min with a flow rate of 250 nl/min. The peptides were ionized by electrospray ionization at +2.0 kV. Tandem mass spectrometry analysis was carried out on a Q-Exactive Plus mass spectrometer using HCD fragmentation. The data-dependent acquisition method acquired MS/MS spectra of the top 10 most abundant ions with charged state ≥2 at any one point during the gradient. MS/MS spectra were searched against the Swissprot reference proteome using Mascot search engine (Version 2.7, Matrix Science) or against human ITGB2 protein sequence using Byonic^TM^ (Version 3.0, Protein Metrics). Precursor mass tolerance and fragment tolerance were set at 10 ppm and the precursor ion charge state to 2+, 3+ and 4+. Variable modifications were defined as oxidized Met, deamidated Asn/Gln, N-terminal pyro Glu/Gln, iodoacetanilide derivative Cys and iodoacetanilide-13C derivative Cys with full trypsin and chymotrypsin cleavage of up to three missed cleavages. Only peptides with a peptide score >30 (p<0.05) and error <6 ppm were selected for quantification of relative abundance (**Supplementary File 1 Table S2**). Relative ion abundance of peptides labelled with ^12^C-IPA and/or ^13^C-IPA in extracted ion chromatograms generated using XCalibur Qual Browser software (Thermo Fisher Scientific, Waltham, Massachusetts, v2.1.0). The redox state of cysteine was calculated as a percentage of the abundance of ^12^C-IPA labelled peptide of the total sum of abundance of ^12^C-IPA and ^13^-IPA labelled peptide.

The mass spectrometry proteomics data have been deposited to the ProteomeXchange Consortium via the PRIDE (Perez-Riverol et al., 2022) partner repository with the dataset identifier PXD032688. The dataset is currently private but is accessible using the following login details at https://www.ebi.ac.uk/pride/login.

Username

Password: F828dFcq

### Recombinant expression of Mac-1 on BHK cells

cDNA constructs for expression of recombinant Mac-1 in mammalian cells were generated by GenScript. *ITGAM cDNA* was cloned into the vector pcDNA3.1/HygroB(+). Wild type (WT) *ITGB2* cDNA or Cys to Ser mutant DNA (C169SC176S, C224SC264S, or both) was cloned into the vector pcDNA3.1/Neo(+). cDNA was cloned using the restriction enzyme sites HindIII and Xbal.

*ITGAM* and *ITGB2* cDNA constructs were linearized with restriction enzymes SspI and Fsp1 (New England Biolabs) respectively and were cotransfected into BHK cells using Lipofectamine 2000 (Thermofisher) according to manufacturer’s protocol. Stably transfected cells were selected by culturing in media containing hygromycin B and G418 (Thermofisher); cells were cultured in DMEM supplemented with L-glutamine at 2 mM, 10% v/v fetal calf serum, hygromycin B at 500 µg/mL, and G418 at 500 µg/mL. Anti-CD18 antibody H52 (Developmental Studies Hybridoma Bank) was conjugated with Alexa Fluor 488 using Alexa Fluor 488 protein labeling kit (ThermoFisher) as described by manufacturer’s instruction. Expression of Mac-1 in transfected BHK cells was then measured by incubating cells with 10 µg/mL of Alexa Fluor 488 conjugated anti-CD18 antibody H52 or APC conjugated anti-CD11b antibody OKM1 (Boster Biological Technology) at 5 μg/mL for 30 min at 25°C, washed, and analyzed by flow cytometry on a BD Accuri C6. Expression was determined by comparing with stained vector BHK cell transfects.

### Static BHK adhesion assays

Wells of a 96-well plate were coated with 10 μg/mL of ICAM-1/Fc at 4°C for 16 h. Wells were then washed with PBS and blocked with 1% w/v polyvinylpyrrolidone for 2 h at room temperature, followed by washing with PBS. 100 μL of BHK cells at 1×10^6^ cells/mL, expressing wild type Mac-1 or disulfide mutants, was then added to each well and left to adhere for 2 h at 37°C. After 3 gentle wash steps with PBS, 100 μL of 1 μg/mL calcein AM was added to each well and left to stain for 30 min at 37°C. The fluorescence of each well was then measured at 488/520 nm ex/em on a Tecan M1000 plate reader.

### Cell de-adhesion assay under flow

Microfluidic devices were produced from PDMS (Dow Corning) and assembled as previously described (Dupuy et al., 2019). Microfluidic devices were coated with 10 µg/mL ICAM-1/Fc for 5 h at 25°C. Devices were then blocked with BSA (1% w/v) for 1 h, then washed with PBS. Recombinant PDI purified from *E. coli* was either reduced with 10 mM DTT in PBS for 30 min at 25°C or oxidized by 200 µM oxidized glutathione (GSSG) in PBS for 16 h at 25°C. PDI was desalted into PBS using zeba desalting columns. BHK cells (1 x 10^6^ cells/mL) were stained with calcein AM (Thermofisher) at 1 µg/mL, perfused without or with 1 µM PDI (reduced or oxidized) through microfluidic devices and left to settle and adhere to surfaces at 25°C for 15 min. Devices were then perfused with PBS for 1 min at a shear stress of 0.175 dynes/cm^2^ (25 s^-1^) to wash off non-adherent cells. Tile scan images were taken on a Zeiss LSM 880 confocal microscope and the shear force was doubled every min until 11.2 dynes/cm^2^ (1600 s^-1^) was reached. Adherent cells were then quantified and normalized to the number of cells adherent at 0.175 dynes/cm^2^.

The area under each curve was calculated in GraphPad Prism 9. Data was also best fitted using nonlinear regression for one phase exponential decay in GraphPad Prism 9 to calculate the decay constant (*K,* cm^2^ dynes^-1^) and shear force (*F_50_*, dynes/cm^2^) at which 50% of BHK cells were de-adhered from ICAM-1.

### Detection of Mac-1 conformation states

Mac-1 expressing BHK cells (1×10^5^ cells) in PBS containing 1 mM CaCl_2_ and 1 mM MgCl_2_ were stained separately with either FITC-conjugated anti-CD18 antibody MEM48 (Thermofisher) at 1:100 dilution or APC-conjugated CBRM1/5 at 0.5 μg/mL, washed, and measured by flow cytometry. Cells were stained with conformation non-specific Alexa Fluor 488 conjugated H52 at 10 μg/mL as a control for Mac-1 expression. Binding of antibodies was calculated as a percentage shift from staining of BHK cells transfected with vector alone. The number of H52+ cells were normalized to the expression ratio of CD11b+ cells (OKM1+) and CD18+ cells (H52+). The distribution of cells in each of the Mac-1 conformations was determined as follows: % Extended open = [% CBRM1/5+ cells]; % Extended closed = [% MEM48+ cells] – [% CBRM1/5+ cells]; % Bent = [% H52+ cells] – [% MEM48+ cells]

### Molecular dynamics simulations

Molecular dynamics (MD) simulations of the integrin β-propeller–βI complex were carried out. Initial coordinates of the complex were taken from the X-ray structure of the Leukocyte integrin ɑLβ2 (PDB id. 5E6U) (Sen and Springer, 2016). Note that the βI domains of Mac-1 and LFA-1 (ɑLβ2) are identical, while the β-propeller units have a 73% sequence similarity (with 41% residues being identical). The β-propeller consisted of the segments 1–122 and 320–591 of the ɑ domain sequence, while the βI domain corresponded to the amino acids 101–344 of the β2 sequence. Four situations were considered: (i) with both C169-C176 and C224-C264 disulfide bonds formed (“wt”), (ii) with C169-C176 bond reduced, (iii) with C224-C264 bond reduced, and (iv) with both disulfide bonds reduced (“both”). The complex (in any of the four forms) was inserted in a dodecahedral simulation box and solvated by ~35165 water molecules. Four Calcium and one Magnesium ions were observed to be bound to the protein in the crystallographic X-ray structure. These ions were considered in the simulation. Surrounding crystallographic water molecules were also considered. Sodium and Chloride ions were added at a concentration of approximately 0.15 M, with an excess of the earlier to ensure an electrically neutral system. The system contained ~116835 atoms in total.

The GROMACS MD package (version 2020.3) was employed (Abraham et al., 2015). The CHARMM36 force-field was used for the protein,(Best et al., 2012) the CHARMM TIP3P model for the water molecules, and default CHARMM parameters for the ions. Electrostatic interactions were computed with the Particle mesh Ewald method (Darden et al., 1995; Essmann et al., 1995). Short-range non-bonded interactions were modelled with a Lennard-Jones potential, within a distance of 1.2 nm. Neighbor searching was carried out by using the Verlet buffer scheme (Pall and Hess, 2013). Bonds involving protein hydrogen atoms were constrained using the LINCS algorithm (Hess et al., 1998). Both angular and bond stretching internal motions of water molecules were also constrained by using SETTLE (Miyamoto and Kollman, 1992). Equations of motion were integrated using the Leap-Frog algorithm at discrete time steps of 2 fs. Temperature was maintained constant at 310 K by using the Nose-Hoover thermostat ((Berendsen et al., 1984; Nose, 1884) for the equilibration steps), using a coupling constant of 1 ps. Pressure was also kept constant at 1 bar by coupling the system to the isotropic Parrinello-Rahman barostat (coupling constant 5 ps) (Parrinella and Rahman, 1981).

Before molecular dynamics, the potential energy of the system was minimized by using the steepest descent method. Subsequently, the solvent was equilibrated around the protein, during 500 ps at constant volume followed by 1000 ps at constant pressure. During these equilibration steps the protein was maintained position-restrained (elastic constant of 1000 kJmol^-1^nm^-2^). For the subsequent production runs the protein restraints were released. N=10 independent simulation replicas were carried out for each system (n=8 when C169-C176 was reduced). The simulation length of each replica varied from 400 ns up to 447 ns, for a total cumulative simulation time of ~4.3 µs (both cysteines oxidized); ~3.4 µs (C169-C176 reduced); ~4.25 µs (C224-C246) reduced, and ~4.3 µs (both cysteines reduced). The total cumulative simulation time was ~16.3 µs. From each replica the first 150 ns were accounted as equilibration and thus discarded from further analysis.

Principal component analysis (PCA), consisting of the calculation and diagonalization of the covariance positional matrix, was carried out to detect global conformational changes of βI (Amadei et al., 1993). The carbon-alpha atoms of βI were considered for this analysis, after rigid-body removal of both translation and rotation of the center of mass of the whole complex. PCA was carried out concatenating the trajectories of all four data sets. Trajectories were then projected onto the first PCA eigenvector (which accounted for 39% of the total C-alpha positional fluctuations of βI). Histograms of the projections are presented in the main text.

The fraction of simulation time C_i,j_ in which the residue pair (i,j) was found in contact was computed using CONAN (Mercadante et al., 2018). A contact was assumed to be established if the residues came closer than 0.35 nm. F_i,j_ was obtained separately for the four different data sets concatenating all replicas corresponding to each set: C_i,j_(wt), C_i,j_(C169-C176), C_i,j_(C224-C246), and C_i,j_(both). Accordingly the change in contact probability was quantified as the difference ΔC_ij_(X) = C_i,j_(X) − C_i,j_(wt), with X=C169-C176, C224-C246, or both. To assess the statistical significance of the change in contacts ΔC_ij_(X), C_ij_ was computed separately for each individual replica: C_ij_^r^ (with r=1,…,N replicas with non-negligible C_ij_^r^ values). Only pairs with r≥5were considered for this analysis. The following normalized difference function was computed z=: [ <C_ij_(X)>_r_ − <C_ij_(wt)>_r_ ] / [ σ^2^_r_ (C_ij_(X)) + σ^2^_r_ (C_ij_(wt) ]^½^, with <>_r_ and σ^2^_r_ denoting average and standard deviation squared over the r replicas, respectively. Residue pairs with z>0.5 were considered (main text) and the dependency on z was monitored by setting z> 0.25, 0.5, 0.75, and 1 (**Figure 5 – figure supplement 1**). In all cases pairs with non-negligible change were selected, meaning |ΔC_ij_|>0.4% of the total simulation time.

The non-bonded pair-wise force F_i,j_ between the residues i and j was extracted from the simulations by using force distribution analysis (version 2.10.2) (Costescu and Grater, 2013). Analogously to the change in contacts, pair-wise force differences, with respect to the fully oxidized system, were computed: ΔF_ij_(X) = <F_i,j_(X)> − <F_i,j_(wt)>, with X=C169-C176, C224-C246, or both, and with <> denoting time-average over the concatenated trajectory. Similarly as with the contacts, the z normalized difference was determined by computing per-replica time-averages of the pair-wise forces. z> 0.25, 0.5, 0.75, and 1 and |ΔF_ij_|>1 pN threshold values were applied.

The root mean square fluctuation (RMSF) of the atomic positions of each residue was computed and the following difference was considered for each system: ΔRMSF=RMSF(X)-RMSF(wt), with X=C169-C176, C224-C246, or both. The solvent accessible surface area (SASA) of the βI domain and some subregions of it (indicated in the main text) was extracted from the simulations. Distributions of this quantity for the different systems are presented in the main text.

PCA, RMSF, and SASA calculations were carried out with the GROMACS gmx tools (Van Der Spoel et al., 2005).

### Neutrophil crawling under flow

PDMS microfluidic devices were coated with 10 μg/mL of ICAM-1/Fc for 2 h at room temperature. Microfluidic devices were blocked with 0.5% BSA for 1 h at room temperature. Neutrophils (1×10^6^ cells/mL) were stained with 1 μg/mL of calcein AM for cell tracking and primed with 1 μM fMLF. Neutrophils were added onto microfluidic devices, allowed to settle and adhere for for 5 min. Adherent neutrophils were then exposed to either 0.7 dynes/cm^2^ (100s^-1^) or 5.6 dynes/cm^2^ (800s^-1^) of shear by perfusing Hank’s buffered saline solution containing 5 μM fMLF. Neutrophil crawling was imaged every second using a 40x oil objective (1.2 NA) on an Olympus IX81 fluorescent microscope.

To measure neutrophil crawling, the centroid of each neutrophil was defined as the cell’s position and was determined by thresholding calcein fluorescence on ImageJ. The cell positions between each frame were then used to determine displacement. The net displacement was calculated by the difference of position at the beginning and end of the time as described by Buffone et al. (Buffone et al., 2018, 2019). Crawling speed (µm/min) was calculated from the total distance traveled by a neutrophil from its initial position as determined by cell tracking and dividing it by the total time of migration. The migration index in X-direction is defined as the ratio of the difference between the initial and final X-displacement over the total distance traveled by a cell (X_end_ − X_initial_)/Distance_total_. When migration is near 0, there is no preferred direction in cell migration. When migration index for X-displacement is near −1, it indicates cell migration against the direction of flow, whereas migration index for X-displacement is near +1, it indicates cell migration in the direction of flow. The migration index in Y-direction is defined as the ratio of the difference between the initial and final Y-displacement over the total distance traveled (Y_end_ – Y_initial_)/Distance_total_. Only single cells that remained in the field of view in the duration of experiment were analyzed. Dividing cells and clusters of cells were excluded from analysis.

### Modelling of Mac-1

To model an extended and open structure of Mac-1, the β2 integrin from αXβ2 (PDB:3K6S) was structurally aligned to the β3 chain from the open structure of αVβ3 (PDB:3IJE) (Xiong et al., 2009), and a model structure of αM was built using the αX structure from PDB:5ES4 (Sen and Springer, 2016) as a template. Disulfide bond C770-C776 in αM was manually broken using PyMOL Molecular Graphics System Version 2.0 Schrödinger, LLC to simulate the opening of the hinge between calf1 and the thigh domains, and disulfide bond C461-C492 in β2 was manually broken to simulate the opening of the hinge between EGF1 and EGF2 domains. Residues 1-770 and 776-1099 from αM were structurally aligned to the residues 1-600 and 661-967 respectively in αV from PDB:3IJE. Residues 1-461 and 469-674 from β2 were structurally aligned to residues 1-488 and 449-695 respectively in β3. The extended Mac-1 model was further refined using Fiberdock server for flexible induced-fit backbone refinement (Mashiach et al., 2010). The extended model was assessed using Swiss-model assessment server (Waterhouse et al., 2018). To reduce atom clashes between the two chains, a structural energy minimization was performed in GROMACS (version 2020.2) using the steepest descent integrator with CHARMM27 forcefield.

### Statistical analysis

Unless otherwise stated, data were analyzed by two-tailed, paired student’s t-test. Multiple data sets were analyzed with one-way ANOVA, with Dunnett’s post-hoc multiple comparisons.

## Data availability statement

The mass spectrometry data is available via ProteomeXchange with the dataset identifier PXD032688. All other data generated or analyzed are included in the manuscript and supporting files, and are also available from the corresponding authors.

## Acknowledgements

The authors thank Profs Bruce Furie and Robert Flaumenhaft for providing plasmid used in bacterial expression of redox inactive PDI, Dr Aster Pijning for assistance in flow cytometry data analysis, and Dr Matthew Graus for advice in PDI and Mac-1 colocalization analysis. A.D. is supported by the Australian Government Research Training Program Scholarship. C.A.S. and F.G. acknowledge the support by the Klaus Tschira Foundation and the Deutsche Forschungsgemeinschaft (DFG, German Research Foundation) under Germany’s Excellence Strategy –2082/1 –390761711. P.J.H. is funded by the National Health and Medical Research Council of Australia (grant numbers 1110219, 1143400 and 1143398 to PH) and a Senior Researcher Grant from the NSW Cardiovascular Research Capacity Program. F.H.P. is funded by a Ramaciotti Foundations Health Investment Grant from the Ramaciotti Foundations, a Ministry of Health New South Wales Cardiovascular Early Mid Career Research Grant and a Sydney Cardiovascular Fellowship from the University of Sydney and Heart Research Institute. J.C. is funded by the Robert and Helen Ellis Postdoctoral Fellowship and the Tony Basten Postdoctoral Fellowship from the Sydney Medical School Foundation, University of Sydney.

## Authorship Contributions

P.J.H., F.H.P and J.C. conceived the project. A.D. and F.H.P. designed, performed and analyzed data from microfluidic and imaging studies. A.D. and J.C. designed, performed and analyzed data from mass spectrometry. A.Y. built the structural model of extended Mac-1. C.A.S. and F.G. designed, performed and analyzed data from molecular dynamics simulations. All authors contributed to the writing of the manuscript.

## Figure Legends

**Figure 1 – figure supplement 1.**
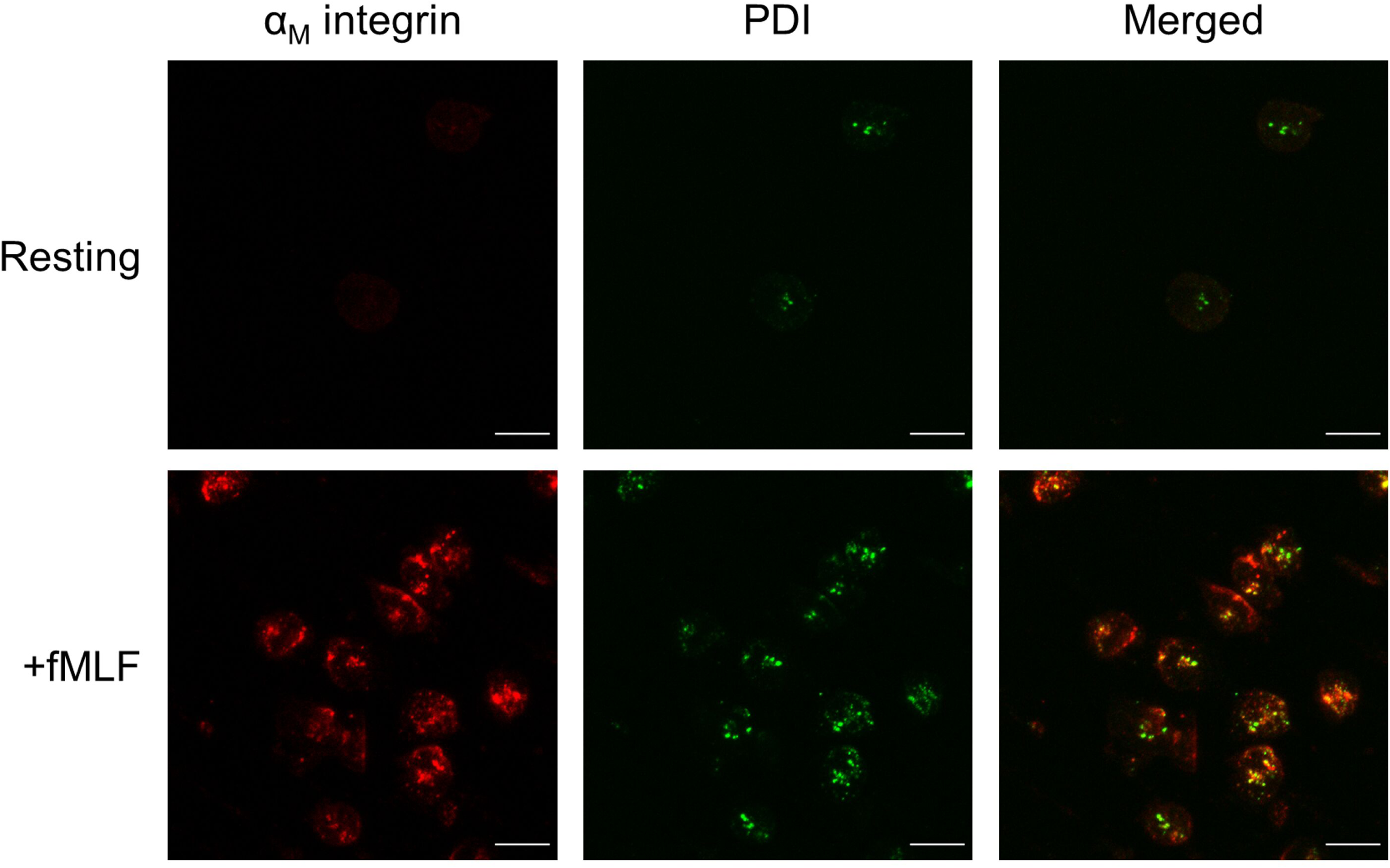
Surface PDI colocalizes with active Mac-1 on neutrophils adhered to ICAM-1 in static condition. Neutrophils were isolated from human blood and stimulated with fMLF. Surface PDI was detected using Alexa fluor 488 conjugated anti-PDI antibody DL-11 (green) on resting or fMLF-stimulated human neutrophils. Active Mac-1 was detected using an APC conjugated anti-CD11b antibody CBRM1/5 (red). After staining with antibodies, neutrophils were added to ICAM-1 coated surface and allowed to adhere before fixing with 4% paraformaldehyde and imaged by confocal microscopy. Scale bar represents 10 µm.

**Figure 2 – figure supplement 1.**
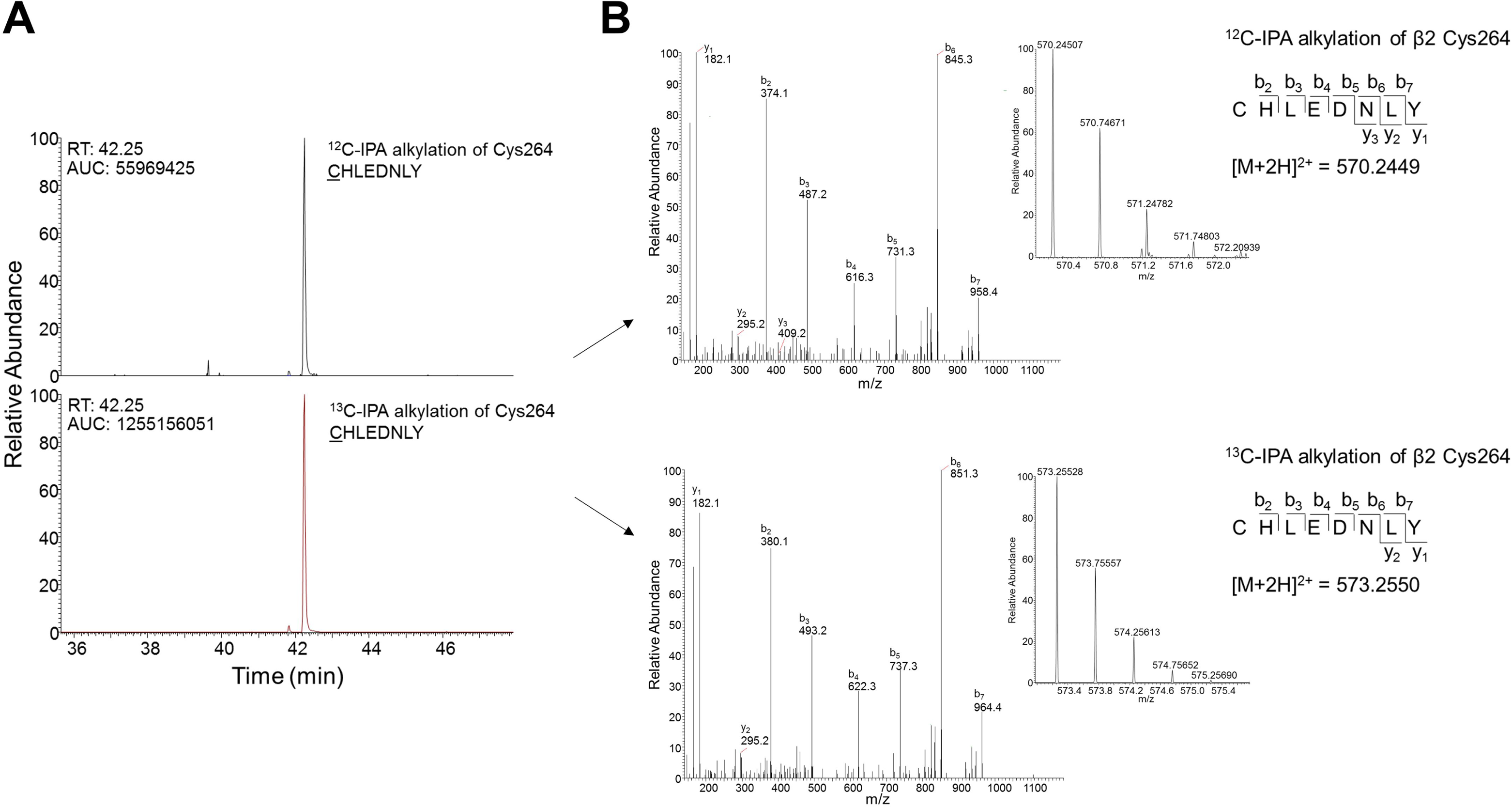
Differential cysteine alkylation and mass spectrometry analysis of cysteine redox state in the β2 integrin. A) Resolution of βI-domain Cys264 containing peptide <u>C</u>HLEDNY with ^12^C-IPA (upper trace) or ^13^C-IPA (lower trace) alkylation under HPLC. B) Representative tandem mass spectra of the CHLEDNY peptide, showing ^12^C-IPA (upper trace) or ^13^C-IPA (lower trace) alkylation of Cys264. The accurate mass spectrum of the peptide is shown in the insets (upper trace, observed [M+2H]^2+^ = 570.2451 m/z and expected [M+2H]^2+^ = 570.2449 m/z; lower trace, observed [M+2H]^2+^ = 573.2553 m/z and expected [M+2H]^2+^ = 573.2550).

**Figure 3 – figure supplement 1.**
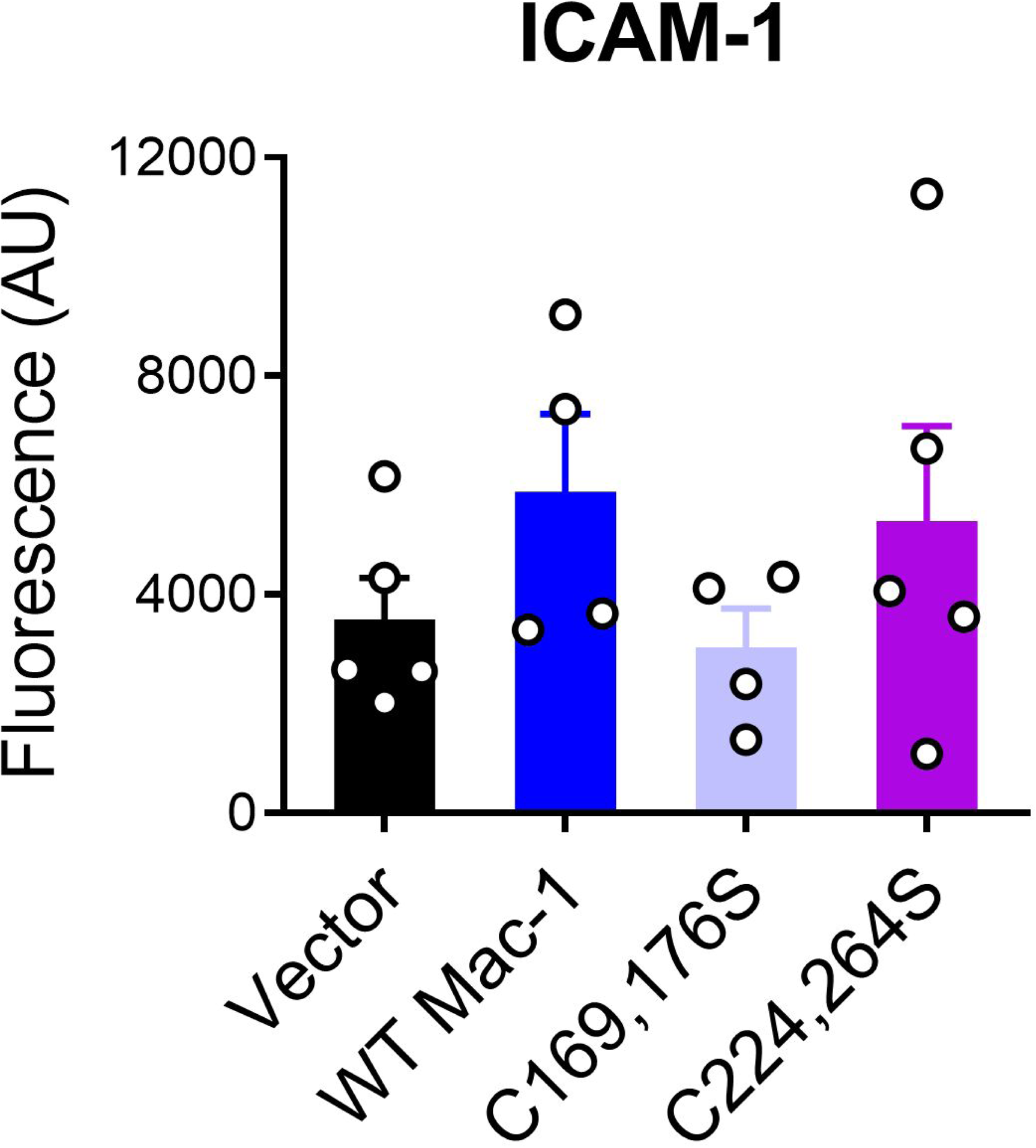
Adhesion of Mac-1 expressing BHK cells to ICAM-1 under static conditions. BHK cells expressing wild type Mac-1 or disulfide mutants were left to adhere on an ICAM-1/Fc coated 96 well plate at 37°C for 2 h. Cells were washed, stained with calcein AM, and fluorescence at 488/520 nm measured. Fluorescence intensity is shown as the mean ± SEM of 4-5 independent experiments.

**Figure 3 – figure supplement 2.**
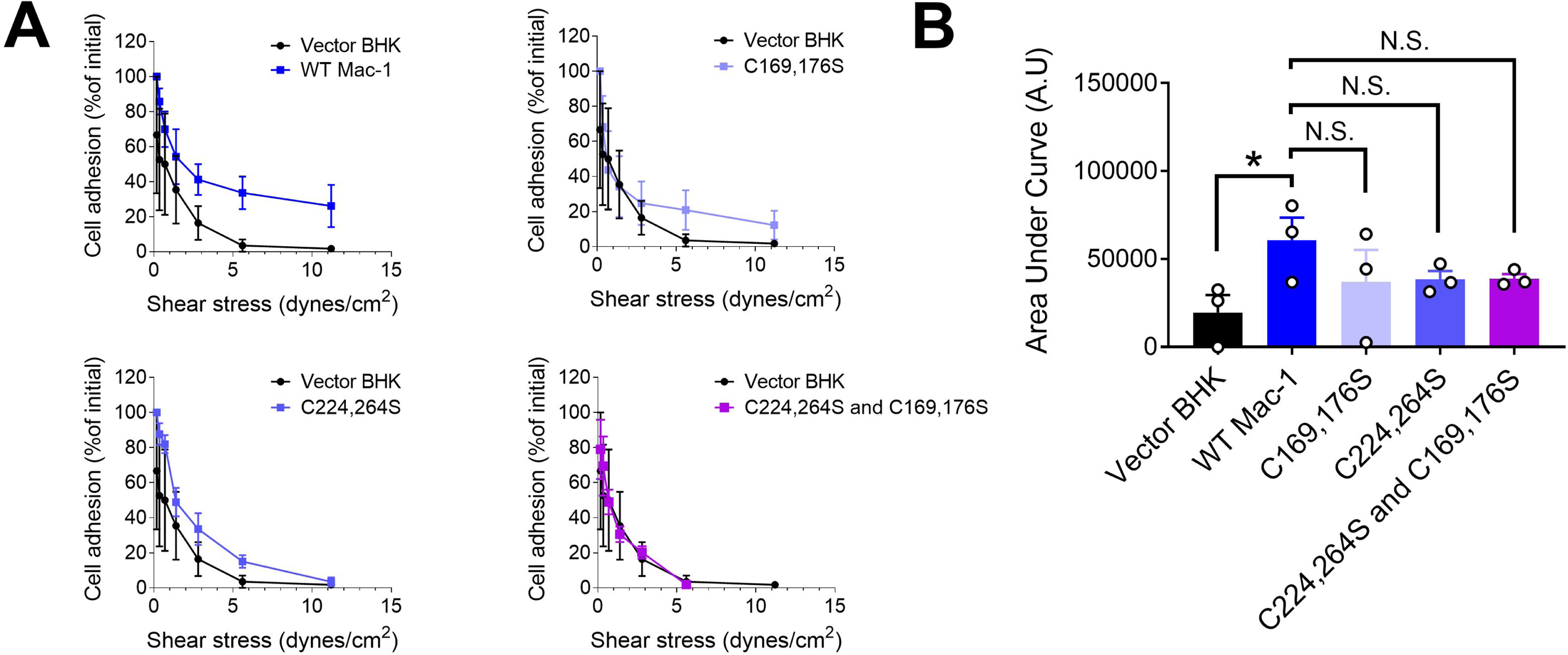
De-adhesion assays of Mac-1 expressing BHK cells from immobilized fibrinogen under increasing shear force. **(A)** Calcein-stained BHK cells were perfused and allowed to adhere to fibrinogen coated microfluidic channel for 15 min. Cells were subjected to shear force at each defined shear rate for 1 min (0.175, 0.35, 0.7, 1.4, 2.8, 5.6 and 11.2 dynes/cm^2^) to allow cell de-adhesion. Images were acquired and the number of adhered cells remained at each shear rate was quantified. **(B)** Area under each curve was from panel B. Data represent mean ± SEM of three biological replicates. P<0.05; N.S.=non-significant by one-way ANOVA with Dunnett’s post-hoc multiple comparisons.

**Figure 4 – figure supplement 1.**
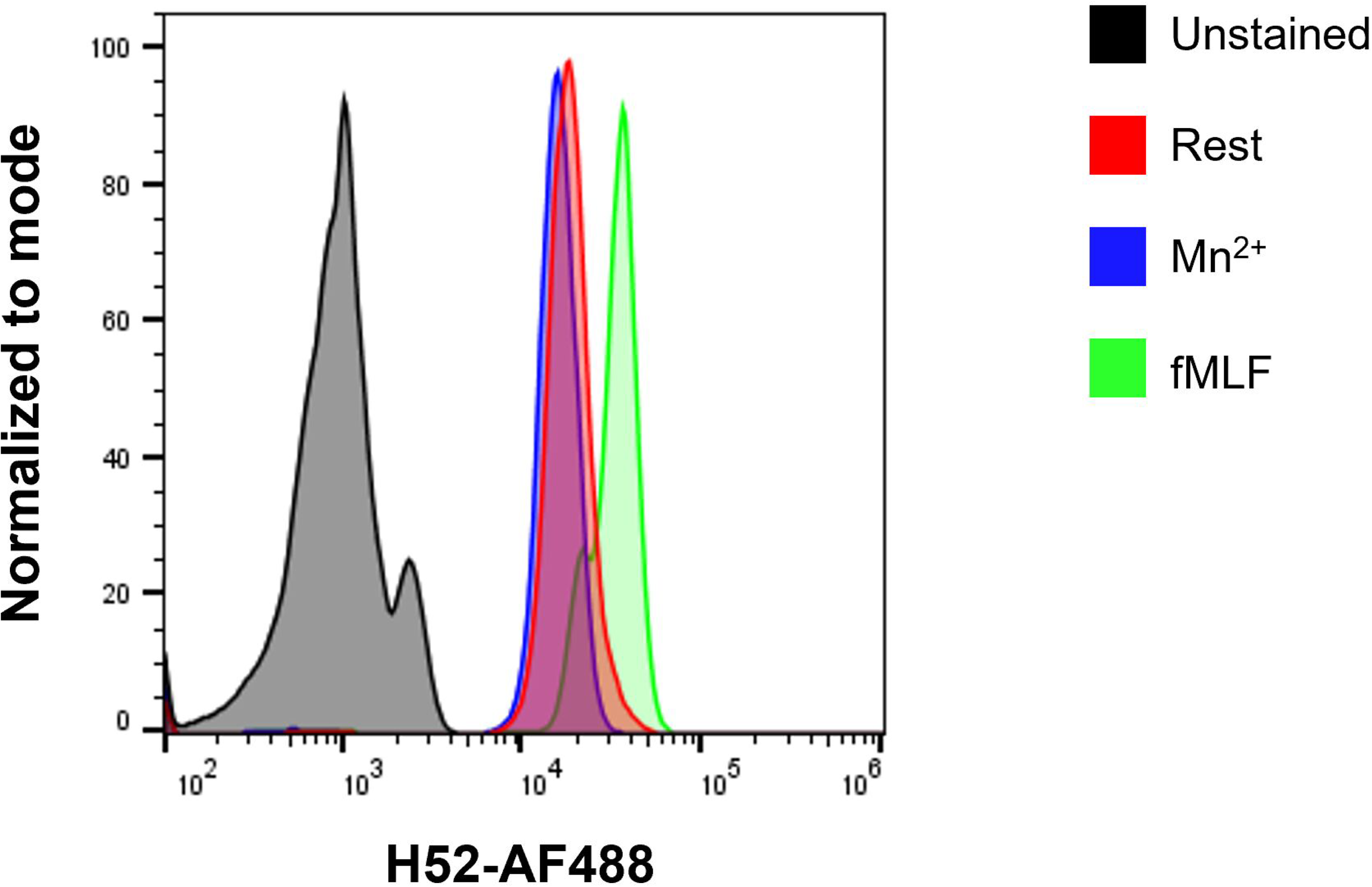
Characterization of pan-conformation anti-CD18 antibody H52. Isolated neutrophils (1×10^6^ cells in 100 µL of HBSS) were incubated with Alexa Fluor 488 conjugated anti-CD18 antibody H52 at 1 µg/mL in the presence or absence of 1 mM MnCl_2_ or 10 µM fMLF for 30 min at 37°C. Samples were then read immediately by flow cytometry on a BD Accuri C6 flow cytometer. The binding of H52 antibody to β2 integrins on the surface of isolated neutrophils is shown at resting (red), upon integrin extension induced by incubation with Mn^2+^ (blue), and upon upregulation of β2 integrins induced by fMLF-stimulation (green) compared to unstained neutrophils (black).

**Figure 5 – figure supplement 1.**
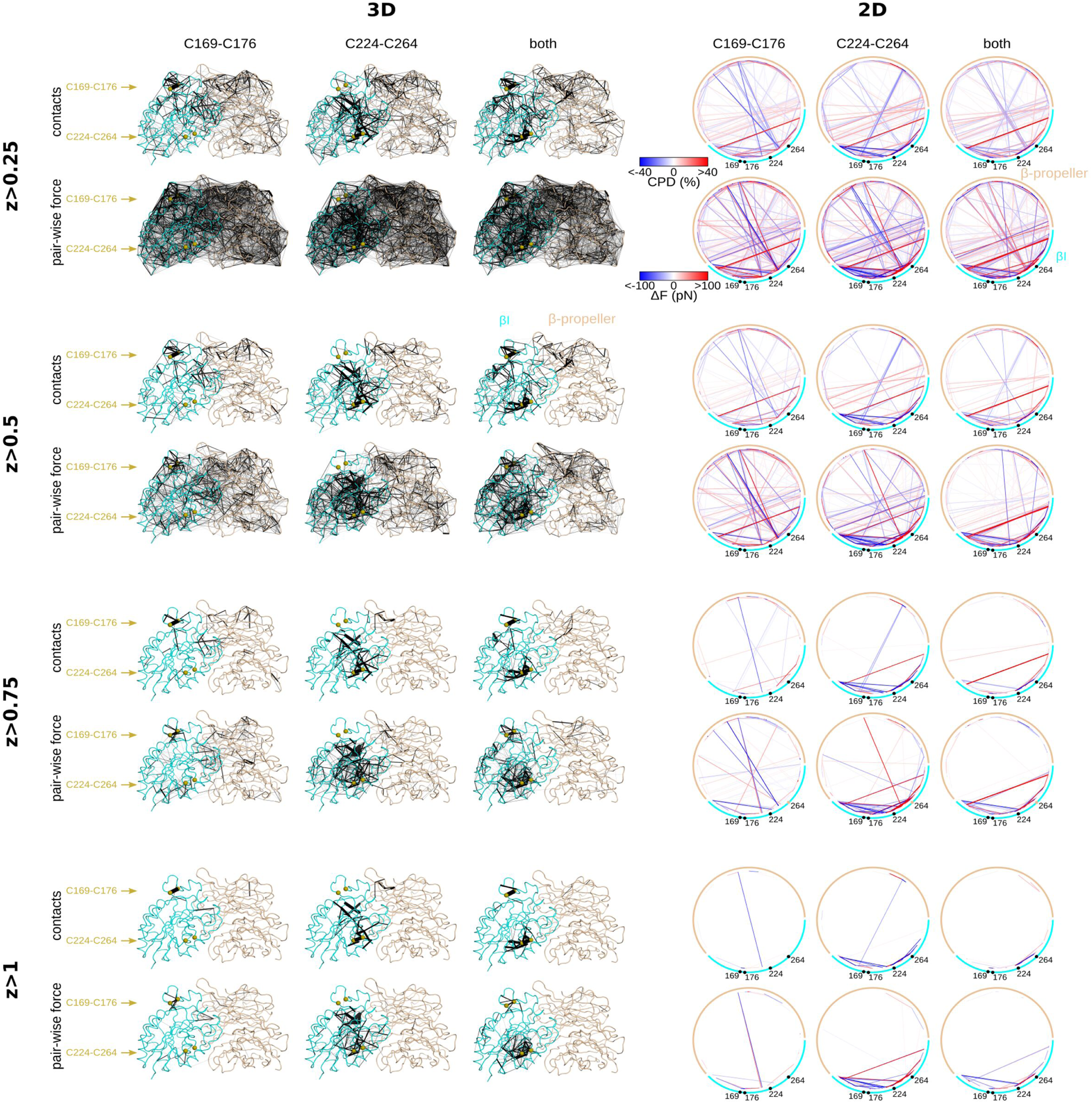
MD simulations. Changes in the pair-wise contact probability ΔC_ij_ and pair-wise force ΔF_ij_ is mapped on the 3D structure (left) and the 2D sequence (right) of the β-propeller–βI domain complex. The amino acid sequence at the right is depicted by a circle where the amino acid is position is given by an angle from 0° (N-terminus of the β-propeller) to 360° (C-terminus of the βI domain). The changes are represented by lines connecting the c-alpha atoms (left) or the points along the circle (right) of residues i and j. The absolute value of ΔC_ij_ or ΔF_ij_ is represented by the thickness of the line (left) or the color according to the shown color bars (right). Same range of values is shown both (as in the color scale) is used both at the left and at the right. Changes are presented for the system, X, with the C169-C176 bond (left column), the C224-C264 bond (middle column) bond, or both bonds (right column) reduced, with respect to the wild-type situation (wt) in which both disulfide bonds are formed: ΔC_ij_=C_ij_(X)-C_ij_(wt) or ΔF_ij_=<F_ij_(X)>-<F_ij_(wt)>. The changes are presented for different normalized differences, z, See definition of z in the methods.

**Figure 6 – figure supplement 1.**
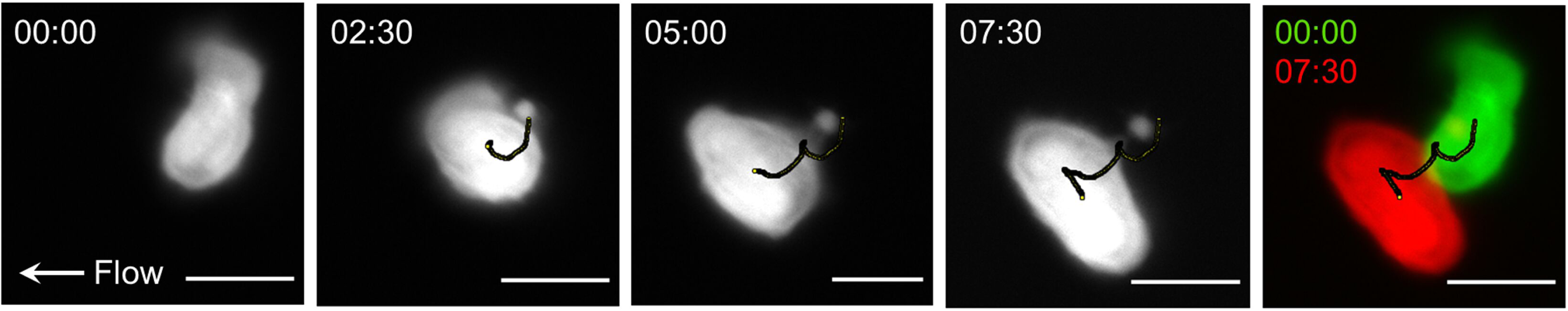
Analysis of neutrophil motility adhered to ICAM-1 under fluid shear. Representative images of the centroid of a single neutrophil used for determining cell track (black line). Initial position (red) and the final position (green) of the cell were measured to determine cell displacement in X- and Y-direction. Time (min) is shown on top right. Scale bar represents 10 µm.

**Figure 6 – figure supplement 2.**
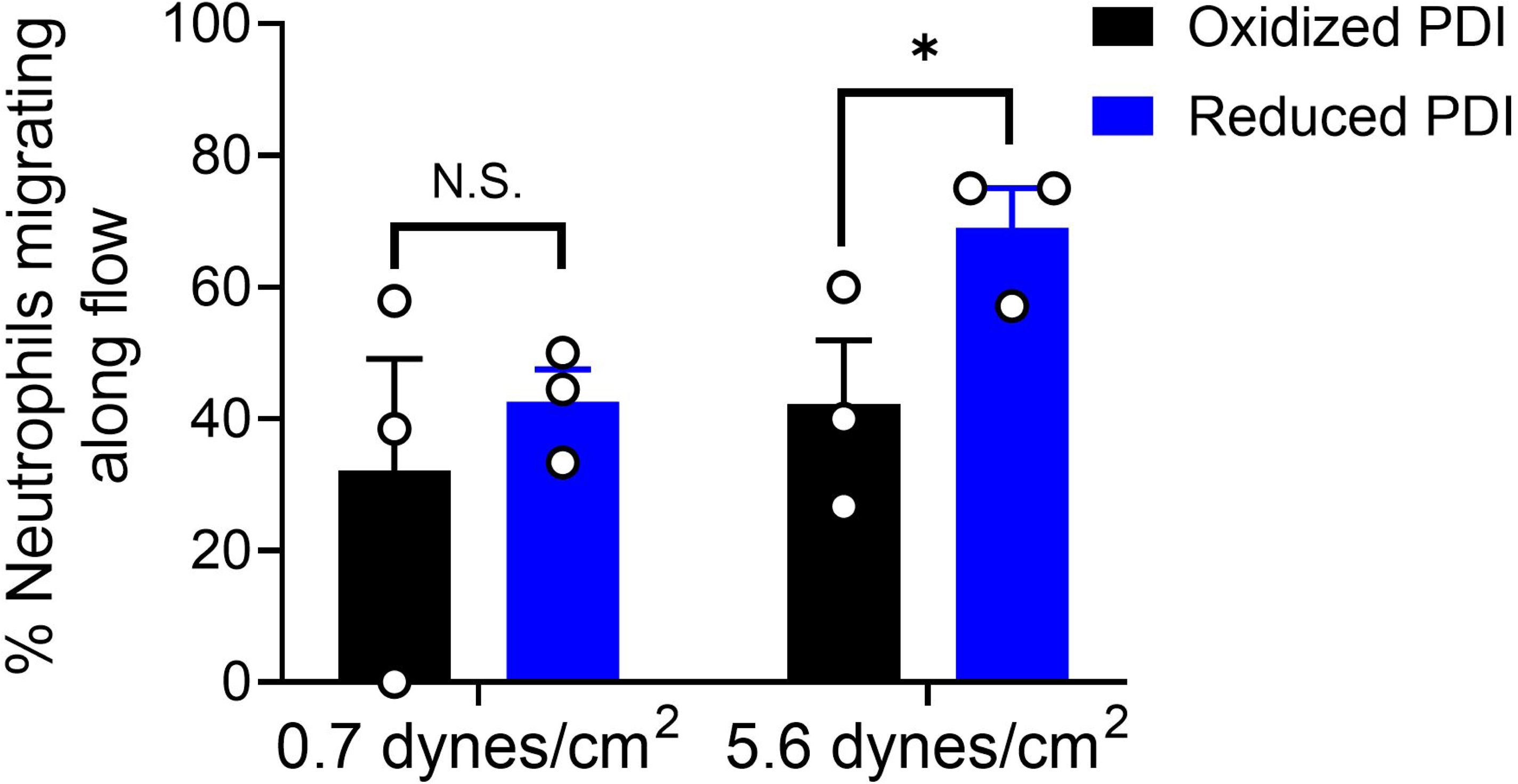
Percentage of neutrophils migration with flow (migration index >0.15). Data shown is mean ± SEM of 3 independent experiments. *P<0.05; N.S. indicates non-significant as assessed by two-tailed, paired student’s ttest.

## Legends for Supplementary Videos 1-4

**Supplementary Video 1.** Crawling of a neutrophil treated with control oxidized PDI on ICAM-1 coated microfluidic chip in presence of 0.7 dynes/cm^2^ fluid shear. Flow direction is from right to left. Scale bar represents 10 µm.

**Supplementary Video 2.** Crawling of a neutrophil treated with reduced PDI on ICAM-1 coated microfluidic chip in presence of 0.7 dynes/cm^2^ fluid shear. Flow direction is from right to left. Scale bar represents 10 µm.

**Supplementary Video 3.** Crawling of a neutrophil treated with control oxidized PDI on ICAM-1 coated microfluidic chip in presence of 5.6 dynes/cm^2^ fluid shear. Flow direction is from right to left. Scale bar represents 10 µm.

**Supplementary Video 4.** Crawling of a neutrophil treated with reduced PDI on ICAM-1 coated microfluidic chip in presence of 5.6 dynes/cm^2^ fluid shear. Flow direction is from right to left. Scale bar represents 10 µm.

